# MEMS directed evolution of two cytochrome P450 enzymes revealing distinct active-sites for convergent functions

**DOI:** 10.1101/2020.12.25.424376

**Authors:** Li Ma, Fengwei Li, Xingwang Zhang, Hui Chen, Qian Huang, Xiaohui Liu, Tianjian Sun, Bo Fang, Kun Liu, Jingfei Chen, Lishan Yao, Dalei Wu, Wei Zhang, Lei Du, Shengying Li

## Abstract

Directed evolution (DE) inspired by natural evolution (NE) has been achieving tremendous successes in protein/enzyme engineering. However, the conventional ‘one-protein-for-one-task’ DE cannot match the ‘multi-proteins-for-multi-tasks’ NE in terms of screening throughput and efficiency, thus often failing to meet the fast-growing demands for biocatalysts with desired properties. In this study, we design a novel ‘multi-enzyme-for-multi-substrate’ (MEMS) DE model and establish the proof-of-concept by running a NE-mimicking and higher-throughput screening on the basis of ‘two-P450s-against-seven-substrates’ (2P×7S) in one pot. With the significantly improved throughput and hit-rate, we witness a series of convergent evolution events of the two archetypal cytochrome P450 enzymes (P450 BM3 and P450cam) in laboratory. Further structural analysis of the two functionally convergent P450 variants provide important insights into how distinct active-sites can reach a common catalytic goal.

## Introduction

Ubiquitous cytochrome P450 enzymes (P450s) are a superfamily of heme-thiolate proteins that catalyze the largest variety of oxidative reactions in nature.^1^ Since the natural functional diversity is believed to be an indicator of a protein family’s evolvability, functionally versatile P450s have become a model system of directed evolution (DE).^2–4^ In past decades, significant successes have been achieved in engineering the stability, catalytic efficiency, substrate scope, regio- and stereoselectivity, and even novel abiological activities of P450s using diverse DE strategies.^5–9^ Today, P450s with desired properties are increasingly demanded because of their broad application in bioproduction of diverse natural products and industrial chemicals.^10^ However, the current DE approaches cannot generate sufficient high-quality P450 biocatalysts to meet the fast-growing demands, thus requiring both new DE strategies with a higher screening efficiency and/or hit rate, and more mechanistic understandings for P450 catalysis to guide the rational enzyme engineering.^11–12^

Compared with DE that normally starts with a single parental protein and ends up with a ‘winner’ mutant with desired properties (‘one-protein-for-one-task’) after rounds of screenings (Figure 1A), natural evolution (NE) targets multiple proteins for multiple tasks simultaneously (Figure 1B), and the overall outcome is a fitter organism with a set of mutant proteins that survives better and reproduces faster under given selection pressure due to the better completed tasks. Apparently, the screening throughput/efficiency of the ‘multi-proteins-for-multi-tasks’ NE is higher than the ‘one-protein-for-one-task’ DE. Inspired by this difference, in this study, we designed and tested a novel ‘multi-enzyme-for-multi-substrate’ (MEMS) DE model (Figure 1C) to mimic NE in principle for higher-throughput screening. Outstanding mutants were efficiently obtained through running a pilot MEMS DE approach on the basis of ‘two-P450s-against-seven-substrates’ (2P×7S). Further structural analysis of the two selected isoenzymes provide new insights into P450 substrate binding/orientation mechanisms resulted from convergent evolution.

**Figure 1.**
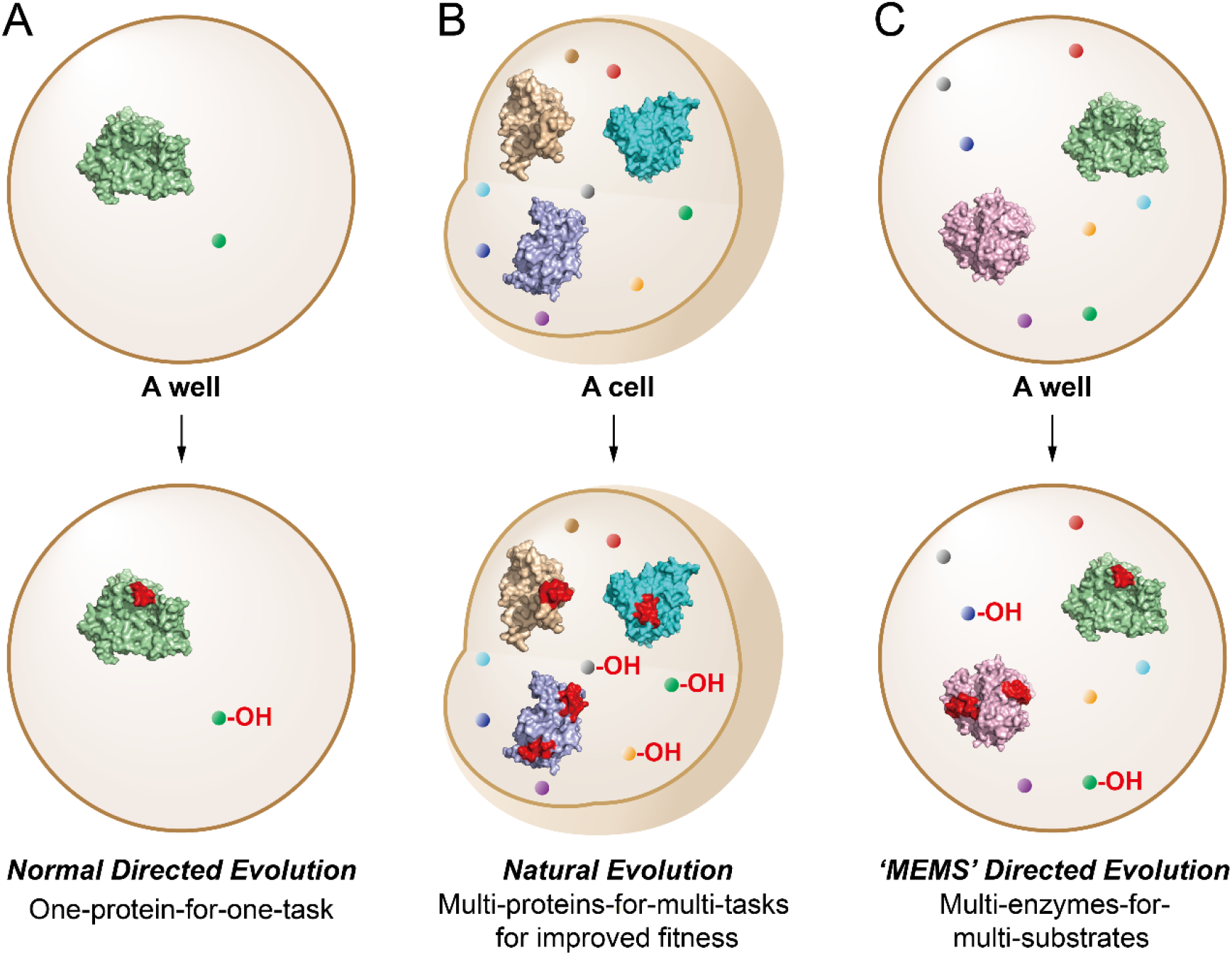
Schematic models of normal directed evolution (**A**), natural evolution (**B**), and ‘MEMS’ directed evolution (**C**).

P450 BM3 (CYP102A1) from *Bacillus megaterium* is a bacterial fatty acid hydroxylase consisting of a P450 domain naturally fused to a mammalian-like diflavin NADPH P450 reductase domain. P450 BM3 is a preferred candidate for P450-based biocatalysis because of its superb catalytic efficiency and self-sufficiency (*i.e*., the P450 activity is independent of separate redox partner proteins).^13^ P450cam (CYP101A1) from *Pseudomonas putida* catalyzes the regio- and stereospecific hydroxylation of (+)-camphor (**1**) to form 5-*exo*-hydroxycamphor (**1a**) when supported by its innate redox partners putidaredoxin (Pdx) and putidaredoxin reductase (PdR)^14^ (Figures S1 and S2). Both enzymes have served as the archetypal models of P450s because of their early discovery, availability of extensive structural information, and the most abundant mechanistic and mutational studies.^15–20^ Notably, the two P450s only share a 17.2% sequence identity at the protein level (Figure S3) and the root-mean-square deviation (RMSD) of their three-dimensional structures is 11.67 (Figure S4), indicative of their distant evolutionary relationship. Thus, in this study we chose these two intensively studied P450s to build our 2P×7S MEMS DE model, in which the seven substrates were **1**, *R*-(+)-limonene (**2**), geraniol (**3**), (+)-valencene (**4**), (−)-ambroxide (**5**), taxadiene (**6**) and indole (**7**) (Figure 2).

**Figure 2.**
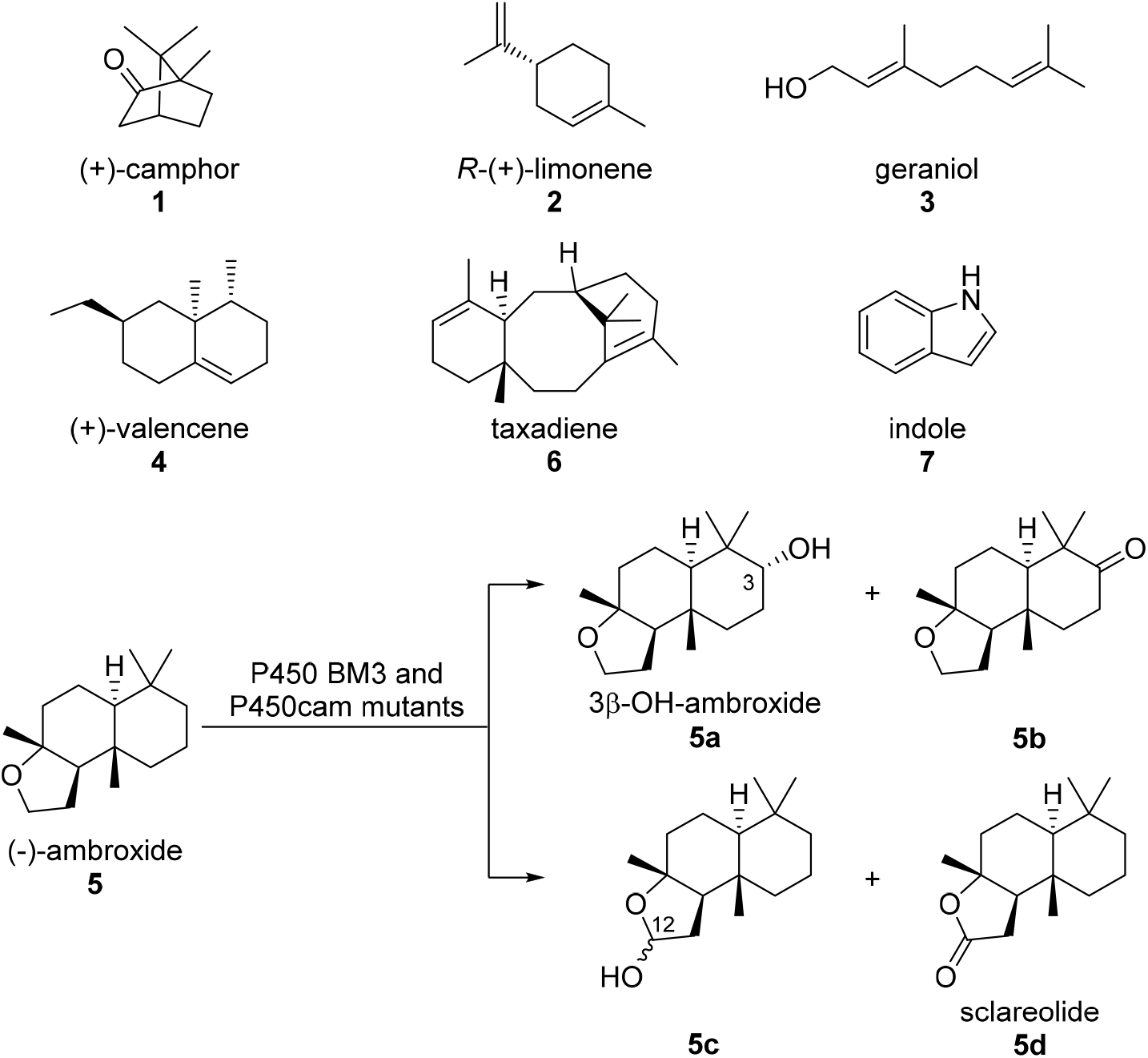
The P450 substrates (**1-7**) used in this study and the oxidative tailoring reactions of (−)-ambroxide (**5**).

To generate P450 mutants for screening, we elected to use the strategy of combinatorial active-site saturation test (CAST)^21^ for both P450 BM3 (the F87A mutant of P450 BM3 with an expanded active site and hence substrate scope was actually used as the starting enzyme;^22^ this mutant will be called P450 BM3 from this point for convenience unless otherwise specified) and P450cam. Based on analysis of the crystal structures of P450 BM3 in complex with a substrate analogue *N*-palmitoylglycine (PDB ID code: 1JPZ)^23^ and P450cam with the native substrate **1** bound (PDB ID code: 2ZWU),^24^ the 11 active-site residues of P450 BM3 were grouped into six sites, including A site (L75 and V78), B site (F81 and A82), C site (A180 and L181), D site (A184 and L188), E site (A328 and A330) and F site (I263); while the 10 active-site residues of P450cam including F87 (A site), Y96 and F98 (B site), L244 and V247 (C site), V295 and D297 (D site), I395 and V396 (E site) and T101 (F site) were selected for CAST (Figure 3A, B). Essentially, all these residues have been identified to be relevant to the activity and selectivity 0f these two P450 enzymes.^11, 25–27^ Grouping the active-site residues into six sites could maximize the cooperative effects with minimal screening efforts. To lower the screening numbers, the NDT codon degeneracy was adopted, which would theoretically give 100% and 85% coverage for the one- and two-residue(s) containing mutation sites, respectively.^28^ Following this design, two P450 libraries, each of which contained 2,016 mutants of P450 BM3 and P450cam, were constructed using the complementary primers (Table S1) that carry the differentially saturated mutations. All these P450 mutants were organized into twelve sublibraries (*i.e*., P450 BM3 A-F and P450cam A-F) according to their mutation sites.

**Figure 3.**
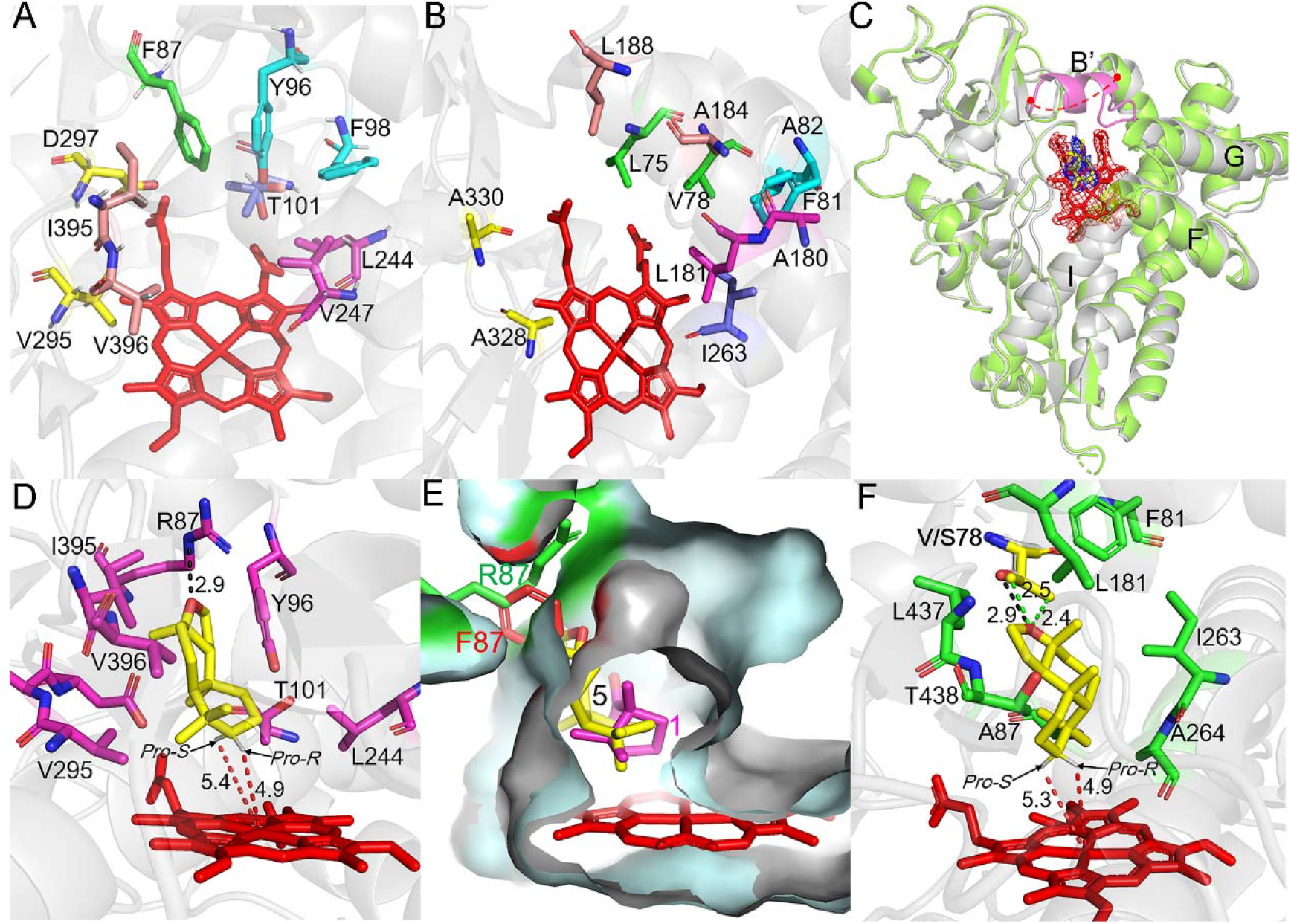
Structural analysis of P450cam and P450 BM3 variants. (**A**) The selected mutational sites of P450cam (PDB ID code: 2ZWU). (**B**) The selected mutational sites of P450 BM3 (PDB ID code: 2X80). The key amino acids grouped into six sites are shown as sticks with different colors in **A** and **B**. (**C**) Superimposition of substrate-free (gray, PDB ID code: 7D4H) and substrate-bound (pale green, PDB ID code: 7D4T) structures of P450cam F87R. B’ helix (90-102 AAs) is absent (shown as the red dashed line representing the missing electron density) and present (colored in magenta) in 7D4H and 7D4T, respectively. The 2Fo-Fc density maps of heme and (−)-ambroxide (**5**) are contoured at 1.0 σ and 0.8 σ, and colored in red and blue, respectively. The substrate **5** is shown as stick in yellow. (**D**) Key residues around **5** within 5 Å shown as sticks in magenta. The substrate **5** is shown as stick in yellow. The distances (in angstroms) are indicated by the dashed lines. (**E**) The binding cavity of P450cam F87R for **5** (pale cyan) and of the wild-type P450cam for camphor (**1**, gray, PDB ID code: 2ZWU). (**F**) The ideal Autodock pose of **5** and the key interacting amino acids. The key residues and **5** are shown as sticks in green and yellow respectively. The distances in angstroms are indicated by the dashed lines.

## Results and discussion

In the 96-well plate-based screening, each individual cell lysates of the P450 BM3 and P450cam (that was co-expressed with its redox partners Pdx/PdR) mutants from the corresponding sublibraries were mixed in the same well that pre-contained a solution of seven substrates **1-7** and NAD^+^/NADP^+^/glucose/glucose dehydrogenase (GDH) to supply P450s with recycled reducing equivalents. Upon the reactions of the two distinct P450s with the mixed seven substrates in a single well (rather than only one reaction of a single mutant with one substrate in a normal DE setting), which in principle mimicking the complex natural cellular enzymatic reactions (Figure 1), the reaction mixtures were extracted with ethyl acetate and the extracts were analyzed by gas chromatography (GC) since there lacks a convenient colorimetric or fluorescent method to detect the oxidative products of **1-6**. In this regard, the two colored products of **7** (Figure S1) acted as visible indicators for the varied activities of P450 mutants.

From the 2,016 chromatograms that carried the activity information of 28,224 reactions, any P450-catalyzed conversions could be readily detected by consumption of one or more substrate(s) and generation of corresponding product(s) (Figure S5). For all the mixed reactions that displayed higher substrate conversion rates than those of either starting P450 enzymes, we performed deconvolution to identify the exact paring of the specific mutant enzyme and the reacted substrate(s) in a one-P450-against-one-substrate (1P×1S) manner, and the active reactions were further confirmed by GC-mass spectrometry (GC-MS) analysis. For all the mutant P450s responsible for improved conversions, their mutation sites were determined by DNA sequencing. Of note, under the screening conditions, wild-type P450cam was only able to recognize **1** (as native substrate), and entirely inactive towards **2-7**; while P450 BM3 showed better substrate flexibility since it could oxidize **2-4** to different extent, but was unable to transform **1** and **5-7** (Figure 4 and Table S2).

**Figure 4.**
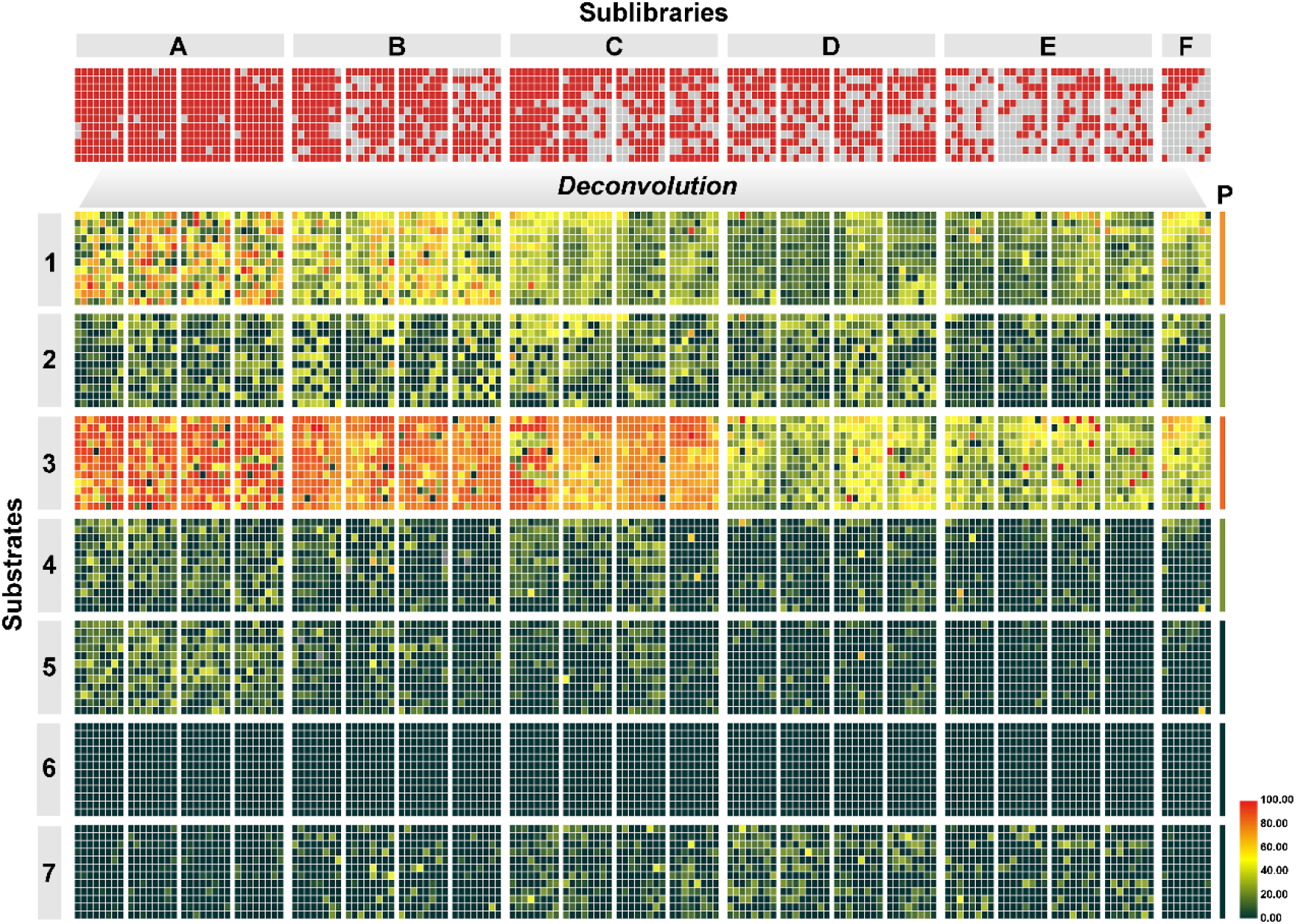
Substrate conversion rates of the whole P450 mutant library. As shown in the *upper* panel, the whole library including 2,016 P450 BM3 mutants of A-F sublibraries and 2,016 P450cam variants of A-F sublibraries were arranged in twenty-one 96-well microtiter plates with each individual well containing one P450 BM3 mutant, one P450cam mutant (together with the co-expressed Pdx/PdR), seven mixed substrates (**1-7**) and NAD^+^/NADP^+^/glucose/GDH. The ‘hit’ and inactive reactions are colored in red and grey, respectively. In the *lower* panel, the GC analytic data for the 2,016 mixed reaction samples were devonvoluted to 28,224 reactivity results of the individually mixed P450 BM3 and P450cam mutants. Two parallel reactions catalyzed by the two mixed parental P450s under the same reaction conditions were carried out as controls. The substrate conversion rates of the control reactions catalyzed by parental enzymes are shown separately in the rightmost panel marked by ‘P’. The color codes in the heat map indicate the differential substrate conversion ratios.

Compared to the normal 1P×1S DE, the 2P×7S DE approach enhanced the throughput of screening for 14 times in a given time. Complete GC analysis of 2,016 samples containing 28,224 reactions (Figure 4) revealed an extraordinarily high hit rate of 68.1% (a well with any reaction showing a higher substrate conversion ratio than either P450 BM3 or wild-type P450cam was defined as a ‘hit’) and distinct reactivity patterns towards different substrates. For instance, none of the tested mutants showed any activity against **6**; while many P450 variants in sublibraries A, B and C demonstrated increased activities towards **3** (see Table S2 for detailed data). In particular, several mutants from sublibrary E of P450 BM3, including A328F/A330F, A328F/A330Y, A328N/A330N, A328N/A330Y, A328N/A330C, A328N/A330F and A328N/A330L showed significantly improved selectivity towards 2,3-epoxy-geraniol (**3a**, Figures S6 and S7). Comparatively, the hit rates of P450 BM3 were 100.0%, 100.0%, 10.3% and 99.1% higher than those of P450cam toward **3, 4, 5** and **7**, respectively. The only exception was **1**, towards which P450cam showed a 66.7% higher hit rate than P450 BM3. Regarding the mutation sites, the hits of both P450 BM3 and P450cam mainly belonged to the corresponding sublibraries A-C for substrates **1-5**. For **7**, sublibraries C-E of P450 BM3 generated more hits than other sublibraries (Table S3).

Taken together, these above results indicate that 1) P450 BM3 is probably a better starting P450 template than P450cam for DE in general; 2) the mutation sites should be carefully selected for different substrates (even for those in the same structural class, *e.g*., terpenoids tested in this study); 3) the selection pressure (the total screening number, mutation rate, substrate concentration, *etc*.) needs to be adjusted according to the challenge level of the target reaction, which could be evaluated by the thermodynamic nature of the reaction, the matching/compatibility between the substrate and the active-site predicted by molecular docking and/or the published mutational data, and the new reactivity data collected from more practices of the current ‘MEMS’ DE strategy; and 4) mutagenesis at different sites could enable the same reaction, highlighting that an enzyme could find different solutions for a common catalytic task.

Of particular interest, a number of isoenzymes were evolved from P450 BM3 and P450cam during the 2P×7S DE, demonstrating an intriguing event of convergent evolution in laboratory. These isoenzymes included 1) P450 BM3 A82F and P450cam F81N/A82F for **7** oxidation (Figure S1); and 2) P450 BM3 V78S, L75F/V78I, L75I/V78S and L75N/V78S as well as P450cam F87R, F87K and F87H for 3*β*-hydroxylation of **5** to form 3*β*-OH-ambroxide (**5a**) (see Figures S6, S8 and S9 for structural determination data). Further oxidation of **5a** into the keto product **5b** (Figure S6) to different extent was also observed under the 1P×1S condition. Since the oxidative reaction of **5** showed high regio- and stereoselectivity, we elected to study the mechanisms for how the two distinct P450 active-sites mediate the common selective oxidation of **5**.

The best ambroxide 3*β*-hydroxylase derived from P450 BM3 turned out to be V78S, which converted 94% of **5** into **5a** in 2 h. Unsurprisingly, this mutant showed a higher substrate binding affinity towards **5** (*K_D_* = 65.0 μM) than its parental enzyme (*K_D_* = 230.0 μM) (Figure S10). We also compared the kinetic parameters of these two enzymes, and the *k*_cat_/*K*_m_ values were determined to be 0.3 and 3.5 μM^-1^·min^-1^ for P450 BM3 and the V78S mutant, respectively (Figure S11). Interestingly, all aquired P450 BM3-based ambroxide 3*β*-hydroxylases including V78S, L75F/V78I, L75I/V78S and L75N/V78S contain an amino acid substitution at V78, indicating that this position is crucial for productive binding of **5** (see below for mechanistic analysis). Thus, we performed saturation mutagenesis at V78 and V78S displayed the highest 3*β*-hydroxylation activity towards **5** among the 20 variants (Figure 5A). Of note, the mutant V78G mainly catalyzed consecutive oxidation at the C12 position, giving rise to 12-OH-amboxide (**5c**) and sclareolide (**5d**) (Figure 2, Figures S2, S6, and S12). Moreover, the introduction of an additional amino acid substitution at the position L75 improved the selectivity of 3*β*-hydroxylation since the further oxidation of **5a** to **5b** (Figures S6 and S13) was inhibited probably due to the significantly attenuated binding of **5a** to the mutants containing the L75 modification.

**Figure 5.**
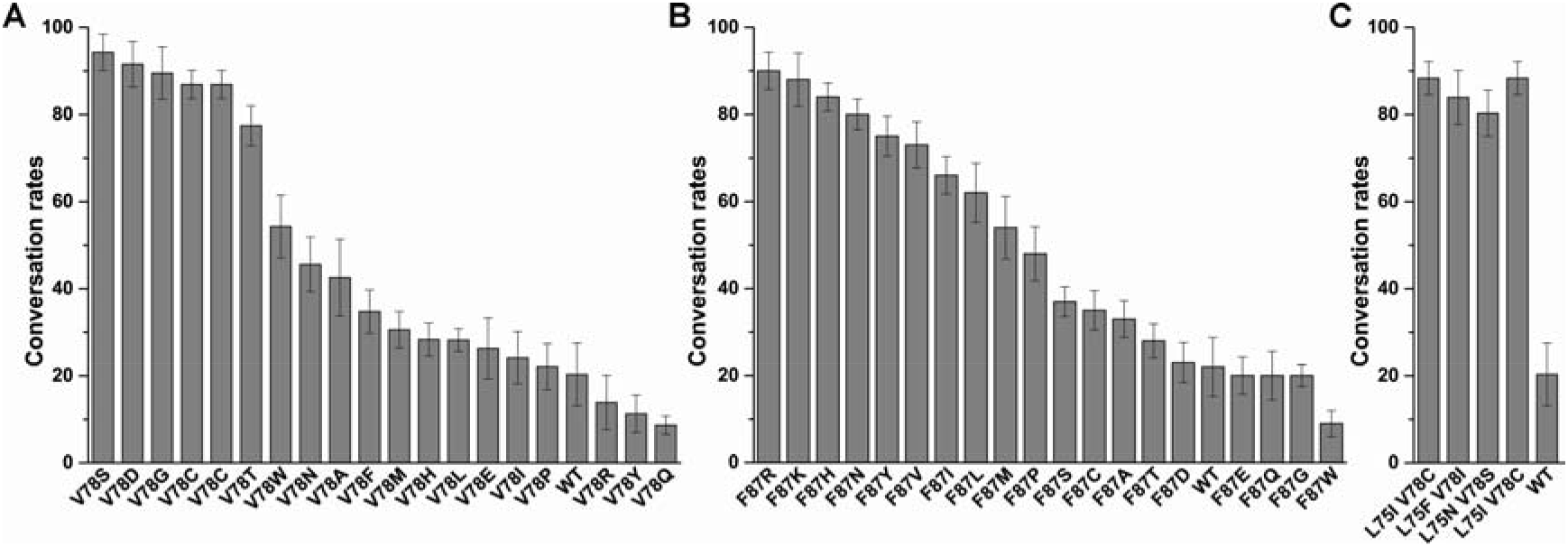
Mutagenesis analysis. **(A)** The conversion rates of **5** by twenty V78 variants of P450 BM3. (**B**) The conversion rates of **5** by twenty F87 variants of P450cam. (**C**) The conversion rates of **5** by three L75/V78 double mutants of P450 BM3. The substrate conversion rates were calculated by (1 - AUC_unreacted substrate_/AUC_total substrate_) × 100% (AUC_total substrate_ was derived from the control reaction using boiled enzyme under identical reaction conditions; AUC: area under curve of the GC ion count chromatograms. All the experiments were carried out in triplicate. The reaction conditions: 0.5 mM substrate **5**, 2 μM P450 enzyme (2 μM Pdx and 2 μM PdR were supplemented for P450cam), and 500 μM NADH (for P450cam) or 500 μM NADPH (for P450 BM3), at 30 °C for 2 h

For P450cam-based ambroxide 3*β*-hydroxylases, we analyzed the most active variant F87R, whose substrate conversion rate was 68% higher than that of wild-type P450cam. The *K_D_* value of P450cam F87R was determined to be 87.3 μM, only slightly lower than that of wild-type enzyme (*K_D_* = 91.1 μM) (Figure S10). Therefore, the increased activity likely resulted from the significantly enhanced *k*_cat_ value, which led to a 3.2-fold increase of *k*_cat_/*K*_m_ (Figure S11). Further saturation mutagenesis analysis showed that the first three places mutants were F87R, F87K and F87H (Figure 5B), which is consistent with the screening results, further supporting the validity of our new ‘MEMS’ DE strategy.

To understand how P450cam F87R and P450 BM3 V78S share the common substrate specificity and catalytic selectivity, we solved the crystal structures of substrate-free P450cam F87R (PDB ID code: 7D4H), **5**-bound P450cam F87R (PDB ID code: 7D4T) and substrate-free P450 BM3 V78S (PDB ID code: 7D4W, only the P450 domain was crystallized, see Supporting Information) at 1.58 Å, 2.1 Å and 2.29 Å resolution, respectively (Figure 3C-F, Table S4). Despite our greatest efforts, we were unable to obtain the structure of P450 BM3 V78S in complex with **5**, possibly due to greasy substrate binding resulted from the spacious substrate entry/egress tunnel and binding pocket (Figure S14).

The substrate-free and substrate-bound structures of P450cam F87R have one and two molecules in an asymmetric unit (Figure S15), respectively. Similar to the previous report,^29^ the electron density of B’ helix (90-102 AAs) in the substrate-free structure is missing in an open state; while B’ helix becomes ordered in a closed state upon **5** binding (Figure 3C). As expected, non-specific hydrophobic contacts dominate the substrate-P450 interactions. Replacement of F87 with an arginine leads to the formation of a key hydrogen bond with the only oxygen atom in **5** (Figure 3D, Figure S16A). The specific anchoring-like interaction determines the substrate position/orientation in the active-site, thereby presenting the *pro*-3*R* C-H bond to be hydroxylated towards the heme-iron reactive center. This observation well explains the regio- and stereoselectivity of 3*β*-hydroxylation by this mutant enzyme. Superimposition of the structure 7D4T and the published P450cam-**1** complex structure (PDB ID code: 2ZWU) further revealed that arginine at position 87 provides extra space for the bigger substrate **5** (than **1**) with its side chain pointing away from the heme group (Figure 3E). Notably, the acidic residue D297 interacts with the side-chain positive charge of R87, giving rise to electrostatic attraction. This observation may also explain why F87R/K/H with a basic substitution showed a higher catalytic activity than other mutants (Figure 5A). Supporting this, the three double mutants of F87R/D297A, F87K/D297A and F87H/D297A unanimously lost most of their activities to oxidize **5**.

In the crystal structure of P450 BM3 V78S, eight molecules in an asymmetric unit was observed and their conformations were consistent with one another (Figure S17A). However, different carbon chain-length polyethylene glycols (PEGs) were co-crystalized with these molecules (Figure S17B). Compared with the published structure of P450 BM3 (PDB ID code: 2X80), their whole three-dimensional structures are almost identical with an RMSD of 0.49. Due to unavailability of the complex structure for P450 BM3 V78S and **5**, we used AutoDock 4.2 to explore the binding mode of **5**. The structure of 7D4W was utilized as a template for creating the starting coordinates. In the top 10 lowest energy docking solutions, we found one ideal catalytic conformation, in which the *pro*-3*R* C-H bond to be oxidized is 4.9 Å (*versus* 5.3 Å for the *pro*-3*S* C-H bond) away from the heme-iron reactive center and the orientation of this C-H bond in substrate is consistent with the experimentally observed stereoselectivity (Figure 3F). As observed in the substrate-bound structure of P450cam F87R, **5** mainly through hydrophobic interactions binds to P450 BM3 V78S via F81, A87, L181, I263, A264, L437 and T438 (Figure 3F, Figure S16B). Interestingly, the V-to-S mutation enables a hydrogen bond to be built between the OH group of serine and the heteroatom in **5**. This anchoring strategy is analogous to the one adopted by P450cam F87R (Figure 3D). Moreover, this point mutation appears to provide extra space to accommodate **5**; V78 in P450 BM3 would otherwise cause steric hindrance since the distance between the closest atom of V78 to **5** is only 2.4 Å (Figure 3F).

Taking into account all the P450 BM3- and P450cam-derived ambroxide 3*β*-hydroxylase mutants, the key mutations are all located in B’ helix, which is the most variable secondary-structure element among P450 families and acts as a flexible lid over the substrate binding pocket together with F/G loop.^30^ It has been well-known that B’ helix is essential for substrate specificity determination in many P450s including P450 BM3 and P450cam (the B’ helix of these two prototypical P450s are quite different, Figure S4). Thus, we suggest that the highly variable B’ helix might be an optimal spot of mutagenesis for substrate specificity engineering. Furthermore, for a hydrophobic substrate such as a terpenoid, the entropy gaining might be the major driving force for its binding to the usually hydrophobic substrate binding pocket of a P450 enzyme. However, the polar heteroaom(s) could be essential to determine the binding mode through more specific substrate-enzyme interactions as we observed in the above structural analysis.

## Conclusions

In this work, we established the proof-of-concept of a novel ‘MEMS’ DE model. This simple but effective strategy significantly improved both screening efficiency and hitrate by mimicking NE that is more ‘violent and chaotic’. ‘MEMS’ DE is particularly useful for the screenings depending on ‘slow’ analytic methods such as gas or liquid chromatography and for studying convergent evolution of proteins/enzymes. It is worth noting that the crossed inhibition effect (although not observed in this study) may occur in a ‘MEMS’ DE approach, which is a potential drawback of this methodology. Nonetheless, this undesired effect could be overcome during a robust ‘MEMS’ DE process provided that a substrate can be better recognized by the hit mutant enzyme than its competitive inhibitor. After all, NE of all enzymes is undergoing in such mixture circumstances.

In the pilot practice of ‘MEMS’ DE for co-evolving P450 BM3 and P450cam, we witnessed a series of convergent evolution events in laboratory. Detailed structural and mutagenesis studies on the two efficient and selective ambroxide 3*β*-hydroxylases, namely, P450 BM3 V78S and P450cam F87R, revealed how the two distinct active-sites accommodate and orient the substrate **5** to achieve the same regio- and stereoselectivity. Inspired by the different solutions of isoenzymes for a common P450 reaction, designing an anchoring residue in B’ helix region to lock the heteroatom in a hydrophobic substrate could be developed into a general strategy for P450 enzyme engineering. More importantly, we envision that the broader application of the ‘MEMS’ DE strategy for a larger repertoire of enzymes in the future will not only provide more high-quality enzymes on-demand in a more efficient manner, but also accumulate much more mutagenesis data for rational design of enzymes on the basis of the big data demanding methodologies including machine learning and artificial intelligence.

## Supporting information

Experimental Procedures,Supplemental Table 1-4,and Figure1-17

## Acknowledgements

This work was supported by the National Key Research and Development Program of China (2019YFA0706900 and 2019YFA0905100), National Natural Science Foundation of China (32025001 and 31872729 to S.L., 31600045 and 32071266 to L.M., 31800664 to F.L., 82022066 to W.Z. and 31800041 to L.D.), the Natural Science Foundation of Shandong Province, China (ZR2019ZD20 to S.L., ZR2016CQ05 to L.M. and ZR2019QC009 to K.L.), the Laboratory for Marine Drugs and Bioproducts of Pilot National Laboratory for Marine Science and Technology (Qingdao) (LMDBKF-2019-01), the Tianjin Synthetic Biotechnology Innovation Capability Improvement Project (TSBICIP-KJGG-001), the State Key Laboratory of Bio-organic and Natural Products Chemistry (SKLBNPC18242), and the Fundamental Research Funds of Shandong University (2019GN030). We thank Zhifeng Li, Jingyao Qu, Jing Zhu, and Xiaoju Li from the State Key Laboratory of Microbial Technology of Shandong University for help in GC-MS, HRMS, and X-ray crystallography testing and analysis. We also thank the beam line BL19U1 of Shanghai Synchrotron Radiation Facility (SSRF, Shanghai, China) for X-ray diffraction data collection.

## References

(1) Ortiz de Montellano, P. R., Cytochrome P450: Structure, Mechanism, and Biochemistry. 4^th^ ed.; Springer International Publishing, Switzerland, 2015.

(2) Arnold, F. H.; Wintrode, P. L.; Miyazaki, K.; Gershenson, A. How enzymes adapt: lessons from directed evolution. Trends Biochem. Sci. 2001, 26, 100–106.

(3) Jung, S. T.; Lauchli, R.; Arnold, F.H. Cytochrome P450: taming a wild type enzyme. Curr. Opin. Biotechnol. 2011, 22, 809–817.

(4) Romero, P. A.; Arnold, F.H. Exploring protein fitness landscapes by directed evolution. Nat. Rev. Mol. Cell Bio. 2009, 10, 866–876.

(5) Zhang, R. J. K.; Chen, K.; Huang, X. Y.; Wohlschlager, L.; Renata, H.; Arnold, F.H. Enzymatic assembly of carbon-carbon bonds via iron-catalysed *sp*^3^ C-H functionalization. Nature 2019, 565, 67–72.

(6) Zhou, H. Y.; Wang, B. J.; Wang, F.; Yu, X. J.; Ma, L. X.; Li, A. T.; Reetz, M. T. Chemo- and Regioselective Dihydroxylation of Benzene to Hydroquinone Enabled by Engineered Cytochrome P450 Monooxygenase. Angew. Chem. Int. Ed. 2019, 58, 764–768.

(7) Prier, C. K.; Zhang, R. K.; Buller, A. R.; Brinkmann-Chen, S.; Arnold, F. H. Enantioselective, intermolecular benzylic C-H amination catalysed by an engineered iron-haem enzyme. Nat. Chem. 2017, 9, 629–634.

(8) Guengerich, F. P.; Martin, M. V.; Sohl, C. D.; Cheng, Q. Measurement of cytochrome P450 and NADPH-cytochrome P450 reductase. Nat. Protoc. 2009, 4, 1245–1251.

(9) Xu, H. F.; Liang, W. N.; Ning, L. L.; Jiang, Y. Y.; Yang, W. X.; Wang, C.; Qi, F. F.; Ma, L.; Du, L.; Fourage, L.; Zhou, Y. J. J.; Li, S. Y. Directed Evolution of P450 Fatty Acid Decarboxylases via High-Throughput Screening towards Improved Catalytic Activity. Chemcatchem 2020, 12, 80–84.

(10) Li, Z.; Jiang, Y.; Guengerich, F. P.; Ma, L.; Li, S.; Zhang, W. Engineering cytochrome P450 enzyme systems for biomedical and biotechnological applications. J. Biol. Chem. 2020, 295, 833–849.

(11) Zeymer, C.; Hilvert, D. Directed Evolution of Protein Catalysts. Annu. Rev. Biochem. 2018, 87, 131–157.

(12) Markel, U.; Essani, K. D.; Besirlioglu, V.; Schiffels, J.; Streit, W. R.; Schwaneberg, U. Advances in ultrahigh-throughput screening for directed enzyme evolution. Chem. Soc. Rev. 2020, 49, 233–262.

(13) Narhi, L. O.; Fulco, A. J. Identification and characterization of two functional domains in cytochrome P-450BM-3, a catalytically self-sufficient monooxygenase induced by barbiturates in *Bacillus megaterium*. J. Biol. Chem. 1987, 262, 6683–6690.

(14) Gunsalus, I. C.; Wagner, G. C. Bacterial P-450cam methylene monooxygenase components: cytochrome m, putidaredoxin, and putidaredoxin reductase. Methods Enzymol. 1978, 52, 166–188.

(15) Whitehouse, C. J.; Bell, S. G.; Wong, L. L. P450(BM3) (CYP102A1): connecting the dots. Chem. Soc. Rev. 2012, 41, 1218–1260.

(16) Ahalawat, N.; Mondal, J. Mapping the Substrate Recognition Pathway in Cytochrome P450. J. Am. Chem. Soc. 2018, 140, 17743–17752.

(17) Poulos, T. L.; Finzel, B. C.; Howard, A. J. High-resolution crystal structure of cytochrome P450cam. J. Mol. Biol. 1987, 195, 687–700.

(18) Follmer, A. H.; Mahomed, M.; Goodin, D. B.; Poulos, T. L. Substrate-Dependent Allosteric Regulation in Cytochrome P450cam (CYP101A1). J. Am. Chem. Soc. 2018, 140, 16222–16228.

(19) Myers, W. K.; Lee, Y. T.; Britt, R. D.; Goodin, D. B. The Conformation of P450cam in Complex with Putidaredoxin Is Dependent on Oxidation State. J. Am. Chem. Soc. 2013, 135, 11732–11735.

(20) Wang, B. J.; Li, C. S.; Dubey, K. D.; Shaik, S. Quantum Mechanical/Molecular Mechanical Calculated Reactivity Networks Reveal How Cytochrome P450cam and Its T252A Mutant Select Their Oxidation Pathways. J. Am. Chem. Soc. 2015, 137, 7379–7390.

(21) Reetz, M. T.; Carballeira, J. D. Iterative saturation mutagenesis (ISM) for rapid directed evolution of functional enzymes. Nat. Protoc. 2007, 2, 891–903.

(22) Whitehouse, C. J. C.; Bell, S. G.; Wong, L. L. P450(BM3) (CYP102A1): connecting the dots. Chem. Soc. Rev. 2012, 41, 1218–1260.

(23) Haines, D. C.; Tomchick, D. R.; Machius, M.; Peterson, J. A. Pivotal role of water in the mechanism of P450BM-3. Biochemistry 2001, 40, 13456–13465.

(24) Sakurai, K.; Shimada, H.; Hayashi, T.; Tsukihara, T. Substrate binding induces structural changes in cytochrome P450cam. Acta Crystallogr., Sect. F: Struct. Biol. Cryst. Commun. 2009, 65, 80–83.

(25) Manchester, J. I.; Ornstein, R. L. Rational approach to improving reductive catalysis by cytochrome P450cam. Biochimie 1996, 78, 714–722.

(26) Bell, S. G.; Chen, X.; Xu, F.; Rao, Z.; Wong, L. L. Engineering substrate recognition in catalysis by cytochrorne P450cam. Biochem. Soc. Trans. 2003, 31, 558–562.

(27) Bell, S. G.; Chen, X. H.; Sowden, R. J.; Xu, F.; Williams, J. N.; Wong, L. L.; Rao, Z. H. Molecular recognition in (+)-alpha-pinene oxidation by cytochrome P450(cam). J. Am. Chem. Soc. 2003, 125, 705–714.

(28) Reetz, M. T.; Kahakeaw, D.; Lohmer, R. Addressing the numbers problem in directed evolution. Chemcatchem 2008, 9, 1797–1804.

(29) Qi, F.; Lei, C.; Li, F.; Zhang, X.; Wang, J.; Zhang, W.; Fan, Z.; Li, W.; Tang, G. L. Deciphering the late steps of rifamycin biosynthesis. Nat. Commun. 2018, 9, 2342–2350.

(30) Sherman, D. H.; Li, S. Y.; Yermalitskaya, L. V.; Kim, Y. C.; Smith, J. A.; Waterman, M. R.; Podust, L. M. The structural basis for substrate anchoring, active site selectivity, and product formation by P450 PikC from *Streptomyces venezuelae*. J. Biol. Chem. 2006, 281, 26289–26297.

