## Supplementary material for "MEMS directed evolution of two cytochrome P450 enzymes revealing distinct active-sites for convergent functions": Experimental Procedures,Supplemental Table 1-4,and Figure1-17

#### Reagents

Antibiotics and chemicals were purchased from SolarBio (Beijing, China) and Sigma Aldrich (St. Louis, MO, USA). KOD-Plus Neo DNA Polymerase was obtained from TOYOBO (Osaka, Japan). Restriction enzymes were bought from Takara (Dalian, China). The kits for plasmid extraction and DNA purification were purchased from OMEGA Bio-Tek (Norcross, GA, USA) and Promega (Madison, WI, USA). His-tagged protein purification used Qiagen Ni-NTA resin (Valencia, CA, USA), Millipore Amicon Ultra centrifugal filters (Billerica, MA, USA) and PD-10 desalting columns from GE Healthcare (Piscataway, NJ, USA). Oligonucleotides synthesis and DNA sequencing were conducted by Sangon Biotech (Shanghai, China).

#### General experimental procedures

The UV-visible spectra were taken using a Spectrophotometer Infinite M200 Pro (Tecan Group Ltd., Switzerland). Pre-coated silica gel plates (GF254, Qingdao Marine Chemical Inc., Qingdao, China) were used for TLC analyses. Silica gel (200–300 mesh, Qingdao Marine Chemical Inc., Qingdao, China) was used for column chromatography (CC). Gas chromatography and GC-MS were carried out using Agilent 7980B and 1200 series instruments (Agilent Technologies Inc., Santa Clara, USA) with an Agilent HP-5 column (30 m × 0.25 mm, 2.5 μm). The program used for GC analysis was as follows: the temperature was increased from 50 ℃ to 260 ℃ at 10 ℃ $min^{-1}$, and maintained at 260 ℃ for 8 min. LC-quadrupole time-of-flight mass spectrometry (LC-Q-TOF/MS) analysis was carried out on a maXis UHR-TOF system (Bruker BioSpin GmbH Co., Rheinstetten, Germany) using a Thermo Scientific Hypersil GOLD column (5 μm, 2.1 mm × 100 mm, Thermo Fisher Scientific Inc., Waltham, USA). Nuclear magnetic resonance (NMR) spectra were acquired on a Bruker 600 MHz spectrometer (Bruker BioSpin GmbH Co., Rheinstetten, Germany). NMR data were processed using MestReNova software. The X-ray diffraction datasets were collected at beam line BL19U1 of the Shanghai Synchrotron Radiation Facility (SSRF, Shanghai, China).

#### Construction of expression vectors

A sequence of full-length P450 BM3 was cloned into pET30a between the *Bam*H I and *Hin*d III restriction sites to afford pET30a-*bm3*. The expression vectors for P450cam and its redox partners Pdx/PdR were constructed as described previously[1]. For construction of the pCDFDuet-1-*pdx*-*pdR* co-expression vector, the *pdx* and *pdR* genes bearing corresponding restriction sites were PCR-amplified and sequentially inserted into the *Nde* I-*Bgl* II and *Eco*R I-*Not* I sites of pCDFDuet-1. The P450cam gene was inserted between the *Nde* I and *Hin*d III restriction sites of pET30a to afford pET30a-*cam*. The plasmids pCDFDuet-1-*pdx*-*pdR* and pET30a-*cam* were co-transformed into *E. coli* resulting the redox-sufficient P450cam biocatalysts. The plasmids pET30a-*bm3* and pET30a-*cam* were used as the template for all P450 BM3 and P450cam variants used in this study.

**Library construction**

Site-specific mutagenesis was performed using a modified QuikChange$^{TM}$ mutagenesis protocol. Primers were designed based on the particular amino acid(s) chosen with NDT codon degeneracy (Supplementary Table 1). The P450s with higher catalytic activities than their corresponding parental enzymes were subject to saturation mutagenesis. The mutant libraries were constructed using overlap PCR with KOD-Plus Neo DNA Polymerase (TOYOBO, Japan). PCR products were analyzed on agarose gel by electrophoresis and purified using the Promega Wizard® SV Gel and PCR Clean-Up System (Madison, WI, USA). The recovered PCR products were digested with *Dpn*I at 37 ℃ for 3 h to remove the template DNA, and then directly transformed into the electro-competent *E. coli* BL21(DE3) cells to construct the mutant libraries.

**Screening in 96-well plates**

Single colonies were randomly picked and innoculated into 300 µL of LB medium containing 50 µg/mL kanamycin in a certain number of sterilized 96-deepwell plates. The cultures were grown at 37 ℃, 220 rpm for 12 h. The overnight culture (40 µL) in each well was transferred into a new sterilized 96-deepwell plate pre-containing 400 µL of TB medium (supplemented with 50 µg/mL kanamycin, 1 mM thiamine, and the rare salt solution: 25 µM $FeCl_3 \cdot 6H_2O$, 4 µM $ZnCl_2$, 2 µM $CoCl_2 \cdot 6H_2O$, 2 µM $Na_2MoO_4 \cdot 2H_2O$, 2 µM $CaCl_2$, 3 µM $CuSO_4$ and 2 µM $H_3BO_3$). This plate was incubated at 37 ℃, 220 rpm for 4 h. Then, the P450 enzyme expression was induced by the addition of 0.2 mM isopropyl-*β*-D-thiogalactopyranoside (IPTG) and 0.5 mM 5-aminolevulinic acid (final concentrations) into each individual wells and conducted at 20 ℃, 230 rpm for 20 h. The cells were pelleted by centrifugation at 3,700 *g* for 10 min and stored at -80 ℃ for later use. The freeze–thaw cell pellets were resuspended in 50 mM potassium phosphate buffer (pH 7.4) (200 µL/well) containing 100 mg/L lysozyme, 300 U/mL DNase I and 10% Triton X-100. The 96-well plates were centrifuged and the supernatants were transferred to a new microtiter 96-well plate, to which 2.8 mM of mixed substrates **1-7** (0.4 mM for each substrate), 0.5 mM NADPH and NADH as electron donors for P450 BM3 and P450cam respectively, and 20 mM glucose/5 U GDH as an NAD(P)H regeneration system were added. The plates were incubated at 30 °C for 16 h, after which the reactions were quenched by adding 300 µL of ethyl acetate, and the mixtures were shaken at 37 ℃ for 30 min for organic extraction. The mixtures were centrifuged at 3,700 *g* for 10 min and the organic phases were directly used as samples for GC or GC-MS analysis (HP-5 column).

**Protein expression and purification**

A single colony of transformant was inoculated into LB medium containing 50 mg/L kanamycin. The overnight seed culture was used for 1:100 inoculation of 0.5 L TB medium containing 50 mg/L kanamycin, 1 mM thiamin, and the rare salt solution. The *E. coli* cells were grown at 37 ℃ for 2–3 h until $OD_{600}$ reached 0.4–0.6, at which IPTG was added to a final concentration of 0.2 mM to induce gene expression, and 0.5 mM 5-aminolevulinic acid was supplemented as the heme synthetic precursor. The cells were cultured at 20 ℃ for another 20 h. The culture was centrifuged at 6,000 *g* for 10 min to pellet cells. The following protein purification was carried out by following the previously developed procedure[2,3]. Purified proteins were flash-frozen by liquid nitrogen and stored at −80 °C for later use.

### Enzyme concentration determination

P450 BM3 and P450cam were purified according to the procedures described previously[4-5]. The functional P450 concentrations were determined from CO-reduced difference spectra[6] using the extinction coefficient of $\varepsilon_{450-490}$ = 91,000 $M^{-1}$ $cm^{-1}$. The concentrations of ferredoxin and ferredoxin reductase were determined by measuring the absorbance at the selected wavelengths. The extinction coefficients used for calculation of concentrations were $\varepsilon_{391}$ = 102,000 $M^{-1}$ $cm^{-1}$ for Pdx and $\varepsilon_{455}$ = 10,400 $M^{-1}$ $cm^{-1}$ for PdR[7].

### Determination of substrate affinity

Dissociation constants ($K_D$) for binding of the substrate **5** to wild-type and mutant P450cam and to the P450 BM3 heme domain were determined by UV-visible absorption titrations[8] using 1 μM P450 protein in 50 mM potassium phosphate buffer (pH 7.4) buffer in a 1-cm path length quartz cuvette. Spectra were recorded during the substrate titrations and the overall changes $\Delta A$ ($A_{peak}$ - $A_{trough}$) values were plotted against the substrate concentrations. The $K_D$ values were deduced from non-linear fitting with the Michaelis-Menton equation.

### Steady-state kinetics

The kinetic assays were analyzed by GC using at least three independent measurements. In a typical experiment the concentration of **5** varied from 0.1 to 2.5 mM, and the P450 enzyme was added to a suitable concentration to make the substrate consumption fall into a linear range. All the required cofactors were added to excess. The reactions were initiated by adding NADPH or NADH and stopped by thoroughly mixing with an equal volume of ethyl acetate at 0, 30 and 60 s. The $k_{cat}$ and $K_m$ values were determined by fitting the velocity data to the Michaelis-Menten equation.

### Product quantification

Products **3a**, **5a** and **5b** were isolated from the *in vitro* enzymatic reactions. Compounds were separated by silica gel column using a linear mobile phase gradient of petroleum ether/ethyl acetate and eluted with 13:1 (**3a**), 9:1 (**5a**) and or 10:1 (**5b**), *v/v*. The products were characterized by $^1$H NMR using $CDCl_3$ as solvent on a Bruker Avance III 600 MHz spectrometer with a 5 mm TCI cryoprobe. Chemical properties of **3a**, **5a and 5b** are as follows:

Compound **3a**: colorless oil; $[\alpha]_D$ = -9.09 (*c* 2.6, in $CH_2Cl_2$); HR-ESI-MS *m/z* 171.1378 $[M + H]^+$ (*calc.* 171.1385); $^1$H NMR (600 MHz, $CDCl_3$): δ 5.08 (t, *J* = 6.5 Hz, 1H), 3.83 (dd, *J* = 12.1, 4.2 Hz, 1H), 3.68 (dd, *J* = 12.1, 6.7 Hz, 1H), 2.97 (dd, *J* = 6.6, 4.3 Hz, 1H), 2.08 (m, 2H), 1.68 (s, 3H), 1.71 – 1.65 (m, 1H), 1.61(s, 3H), 1.50 – 1.44 (m, 2H), 1.30 (s, 3H). The $^1$H NMR spectra of **3a** was related to 2,3-epoxy-geraniol[9].

Compound **5a**: white powder; HR-ESI-MS *m/z* 253.2169 $[M + H]^+$ (*calc.* 253.2168), $^1$H NMR (600 MHz, $CDCl_3$): δ 3.94 – 3.89 (m, 1H), 3.83 (q, *J* = 8.2 Hz, 1H), 3.25 (dd, *J* = 11.1, 5.2 Hz, 1H), 1.09 (s, 3H), 1.00 (s, 3H), 0.85 (s, 3H),

0.80 (s, 3H). The configuration of the newly generated hydroxyl group was deduced by comparing the *J* value of H-3 with that of reported *ent*-3-hydroxyambroxide[10].

Compound **5b**: white powder; HR-ESI-MS *m/z* 251.2014 $[M + H]^+$ (*calc.* 251.2011), $^1H$ NMR (600 MHz, $CDCl_3$): $^1H$ NMR (600 MHz, $CDCl_3$) δ 3.97 (dd, *J* = 13.4, 7.7 Hz, 1H), 3.88 (dd, *J* = 16.5, 8.2 Hz, 1H), 2.65 – 2.57 (m, 1H), 2.49 (ddd, *J* = 16.3, 7.5, 3.5 Hz, 1H), 2.05 – 1.98 (m, 1H), 1.86 – 1.78 (m, 3H), 1.77 – 1.72 (m, 1H), 1.66 – 1.59 (m, 2H), 1.56 – 1.43 (m, 3H), 1.16 (s, 3H), 1.14 (s, 3H), 1.08 (s, 3H), 0.98 (s, 3H).

**X-ray Crystallography**

Crystals of P450cam F87R were grown at 18 ℃ using the hanging drop vapor diffusion method and P450 BM3 V78S was handled using the sitting drop method at 18 ℃. The substrate-free crystal screen droplets consisted of a 1:1 (*v/v*) protein at 18-20 mg/mL and the well solution of 1.3 M sodium citrate tribasic dihydrate pH 6.5. P450cam F87R and **5** (hydroxypropyl *β*-cyclodextrin as solvent) were mixed at a molar ratio of 1:5~10 at 18-20 mg/mL and the co-crystallization was carried out in 200 mM sodium acetate tribasic dihydrate, 100 mM sodium cacodylate trihydrate, pH 6.4, 25% PEG8000. P450 BM3 V78S was concentrated to 30 mg/mL and crystallized in an optimized crystallization condition (0.2 M magnesium chloride hexahydrate, 0.1 M Bis-Tris pH 5.5, 17% w/v PEG3350). The crystals were flash-frozen in liquid nitrogen after the addition of glycerol to 20%. The data sets were integrated and scaled with the HKL3000 package[11]. The structures were determined by molecular replacement with the structure of native P450cam (PDB ID code: 2ZWU) or P450 BM3 (PDB ID code: 2X7Y) as the initial search model with the program Phaser.[12] The programs Refmac5 and Coot9 were used for the refinement and model building.[13-14] Ramachandran plots were generated with Coot9. The statistics for data processing and structure refinement are shown in Supplementary Table 3. The structural presentations were prepared using PYMOL (http://www.pymol.org).

**AutoDock**

Substrate **5** was docked into the structure of P450 BM3 V78S (PDB ID code: 7D4W) in the zwitterionic form using AutoDock 4.2[15]. Regents and water molecules were removed before docking. All side chains were set as rigid body and grid spacing was set to 1 Å. Other parameters remained as their default values. The top 10 lowest energy docking poses of substrate **5** from 2,500,000 searching results were selected, among which one ideal catalytic conformation was used for analysis.

### Supplementary Tables

**Table S1.** List of primers for NDT mutant libraries of P450 BM3 and P450cam.

| Name | Sequence (5' to 3') |
|---|---|
| P450 BM3-A-F: | AGTCAAGCGNDTAAATTTNDTCGTGATTTTC |
| P450 BM3-A-R: | AAAATCACGAHNAAATTTAHNCGCTTGACT |
| P450 BM3-B-R: | GTACGTGATNDTNDTGGAGACGGGTTA |
| P450 BM3-B-F: | TAACCCGTCTCCAHNAHNATCACGTAC |
| P450 BM3-C-R: | AGTATGGTCCGTNDTNDTGATGAAGCAATG |
| P450 BM3-C-F: | CATTGCTTCATCAHNAHNACGGACCATACT |
| P450 BM3-D-F: | CTGGATGAANDTATGAACAAGNDTCAGCGA |
| P450 BM3-D-R: | TCGCTGAHNCTTGTTCATAHNTTCATCCAG |
| P450 BM3-E-F: | TTATGGCCAACTNDTCCTNDTTTTTCCCTTAT |
| P450 BM3-E-R: | ATAAGGGAAAAAHNAGGAHNAGTTGGCCATAA |
| P450 BM3-F-F: | ATTATTACATTCTTANDTGCGGGACACGAAACA |
| P450 BM3-F-R: | TGTTTCGTGTCCCGCAHNTAAGAATGTAATAAT |
| P450cam-A-F: | TCCAGCGAGTGCCCGNDTATCCCTCGTGAAGCC |
| P450cam-A-R: | GGCTTCACGAGGGATAHNCGGGCACTCGCTGGA |
| P450cam-B-R: | GCCGGCGAAGCCNDTGACNDTATTCCCACCTCG |
| P450cam-B-F: | CGAGGTGGGAATAHNGTCAHNGGCTTCGCCGGC |
| P450cam-C-R: | ATGTGTGGCNDTTTACTGNDTGGCGGCCTG |
| P450cam-C-F: | CAGGCCGCCAHNCAGTAAAHNGCCACACAT |
| P450cam-D-F: | CGCTTCTCGCTGNDTGCCNDTGGCCGCATCCTC |
| P450cam-D-R: | GAGGATGCGGCCAHNGGCAHNCAGCGAGAAGCG |
| P450cam-E-F: | CACAAGAGCGGCNDTNDTAGCGGCGTGCAG |
| P450cam-E-R: | CTGCACGCCGCTAHNAHNGCCGCTCTTGTG |
| P450cam-F-F: | TACGACTTCATTCCCNDTTCGATGGATCCGCCC |
| P450cam-F-R: | GGGCGGATCCATCGAAHNGGGAATGAAGTCGTA |

**Table S2.** P450 variants in the combined sublibraries A-F that showed better substrate conversions than the parental enzymes towards seven sbustrates. 'P' stands for the substrate conversion rates of the control reactions catalyzed by the mixed parental enzymes.

| Entry | P (%) | A | B | C | D | E | F | Total hits | Hit-rate (%) |
|---|---|---|---|---|---|---|---|---|---|
| 1 | 73.9 | 62 | 11 | 2 | 1 | 3 | 1 | 80 | 4.0 |
| 2 | 28.6 | 48 | 105 | 112 | 89 | 22 | 13 | 389 | 19.3 |
| 3 | 82.8 | 239 | 135 | 65 | 4 | 11 | 1 | 455 | 22.6 |
| 4 | 23.9 | 54 | 31 | 44 | 17 | 5 | 8 | 159 | 7.9 |
| 5 | 0 | 265 | 150 | 164 | 112 | 64 | 24 | 779 | 38.6 |
| 6 | 0 | 0 | 0 | 0 | 0 | 0 | 0 | 0 | 0 |
| 7 | 0 | 72 | 78 | 181 | 183 | 106 | 5 | 625 | 31.0 |

**Table S3.** The mutants in sublibraries A-F with > 3-fold higher activities than the parental enzymes for substrate **1**, **5** and **7** and with improved regioselectivity for **3** and **4**. P450 BM3 A328N/A330N even produced a new product from **3**.

| **Sublibrary** | **Biocatalysts (mutation sites)** | **1** | **3** | **4** | **5** | **7** |
|---|---|---|---|---|---|---|
| A | P450 BM3 (L75/V78) | - | - | - | L75F, V78I | - |
| | P450cam (F87) | - | - | - | F87H | - |
| B | P450 BM3 (F81/A82) | - | - | F81N/A82Y | A82F | A82F |
| | P450cam (Y96/F98) | - | - | - | Y96N/F98N | Y96N |
| C | P450 BM3 (A180/L181) | - | - | - | A180F/L181F | A180I/L181F<br>A180F/L181V<br>A180Y/L181F<br>A180R/L181H |
| D | P450 BM3 (A184/L188) | - | - | A184N/L188N<br>A184N/L188Y<br>A184I/L188N | A184/L188 | A184N/L188N<br>A184I/L188Y<br>A184N/L188H<br>A184N/L188F<br>A184N/L188Y<br>A184N/L188S |
| E | P450 BM3 (A328/A330) | A328N/A330N | A328F/A330F<br>A328F/A330Y<br>A328N/A330N<br>A328N/A330Y<br>A328N/A330C<br>A328N/A330F<br>A328N/A330L | - | A328C/A330V | A328F/A330F<br>A328L/A330I |
| F | P450 BM3 (I263) | - | - | I263G | - | - |

**Table S4.** Data collection and refinement statistics for P450 BM3 V78S and P450cam F87R structures.

| | **P450 BM3 V78S** | **P450cam F87R** | **5-bound P450cam F87R** |
|---|---|---|---|
| **Data collection** | | | |
| Space group | p212121 | p212121 | p21 |
| Cell dimensions | | | |
| *a*, *b*, *c* (Å) | 105.91, 167.87 227.98 | 65.91, 74.06, 92.81 | 63.21, 105.72, 73.72 |
| a, b, g (°) | 90.00, 90.00, 90.00 | 90.00, 90.00, 90.00 | 90.0, 113.994, 90.0 |
| Resolution (Å) | 50.0-2.30 (2.34-2.3) | 50.0-1.58 (1.61-1.58) | 50.0-2.1 (2.14-2.10) |
| *R*sym or *R*merge | 0.079（0.710） | 0.056 (0.616) | 0.056 (0.469 ) |
| *I* / s*I* | 23.4（3.2） | 33.5 (3.5) | 20.5 (2.62) |
| Completeness (%) | 99.5 (99.8) | 99.8 (99.8) | 99.0 (99.5) |
| Redundancy | 12.4 (12.1) | 6.6 (6.6) | 3.4 (3.3) |
| **Refinement** | | | |
| Resolution (Å) | 2.29 | 1.58 | 2.1 |
| No. reflections | 170525 | 59698 | 46080 |
| $R_{work}$ / $R_{free}$ | 0.218/0.249 | 0.187/0.223 | 0.189/0.249 |
| **No. atoms** | | | |
| Protein | 29495 | 3125 | 6426 |
| Ligand | no | no | 34 |
| Water | 1651 | 410 | 523 |
| *B*-factors | | | |
| Protein | 37.84 | 15.48 | 31.41 |
| Ligand | no | no | 44.65 |
| Water | 33.54 | 29.70 | 37.56 |
| **R.m.s. deviations** | | | |
| Bond lengths (Å) | 0.007 | 0.023 | 0.014 |
| Bond angles (°) | 1.432 | 2.188 | 1.647 |
| Ramachandran Favored/Outliers (%) | 95.87/0.00 | 97.86/0.00 | 97.49/0.00 |

Highest-resolution shell is shown in parentheses.

### Supplementary Figures

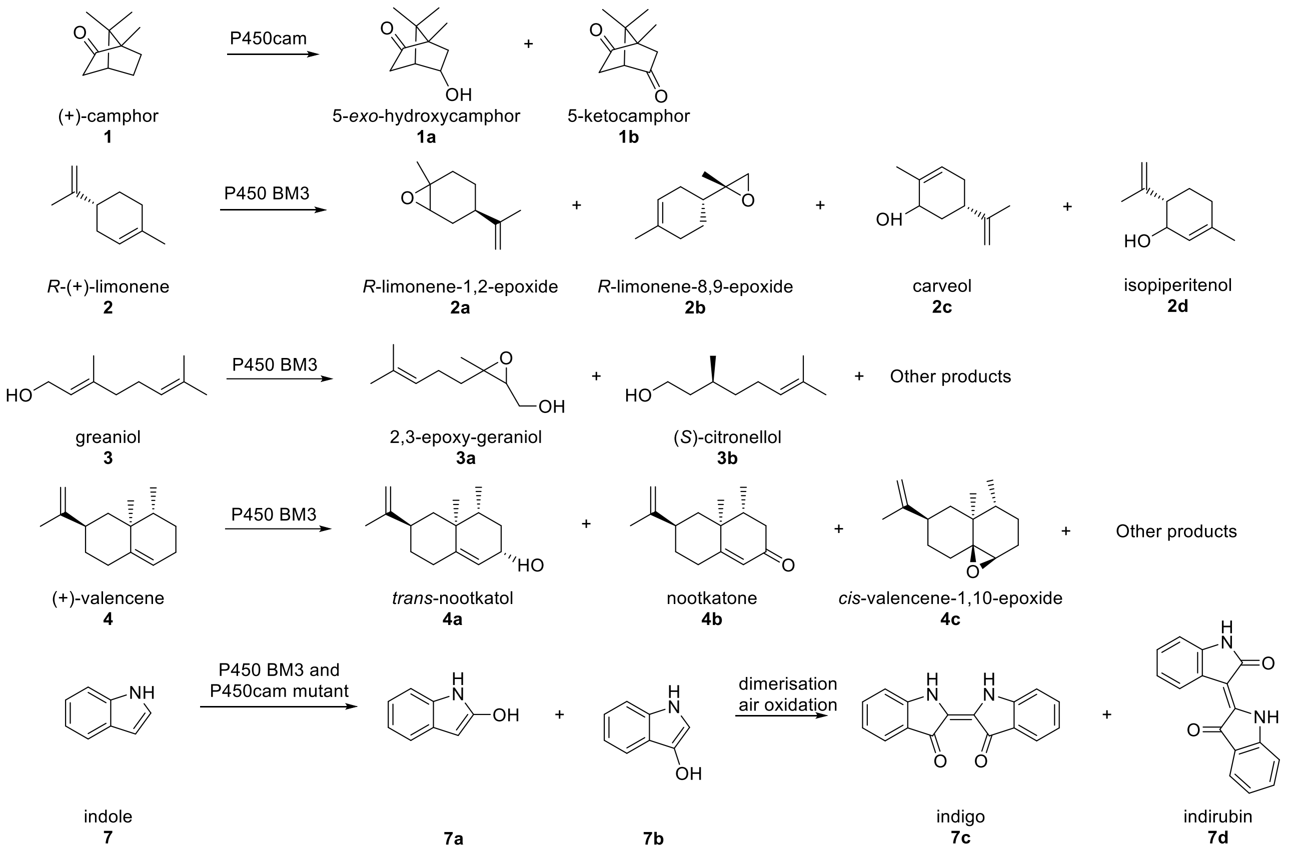

**Figure S1.** The schematic P450 reactions observed in this study. The structures of products were determined by GC-MS and/or NMR.

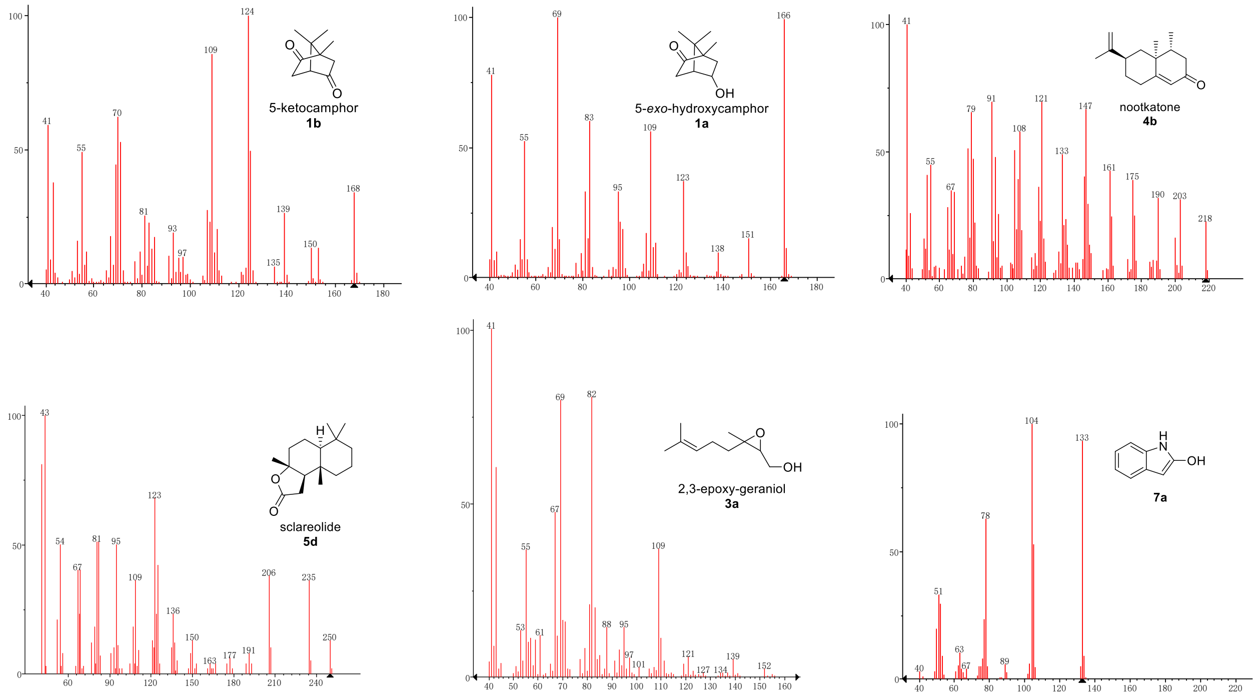

**Figure S2.** GC-MS spectra of products.

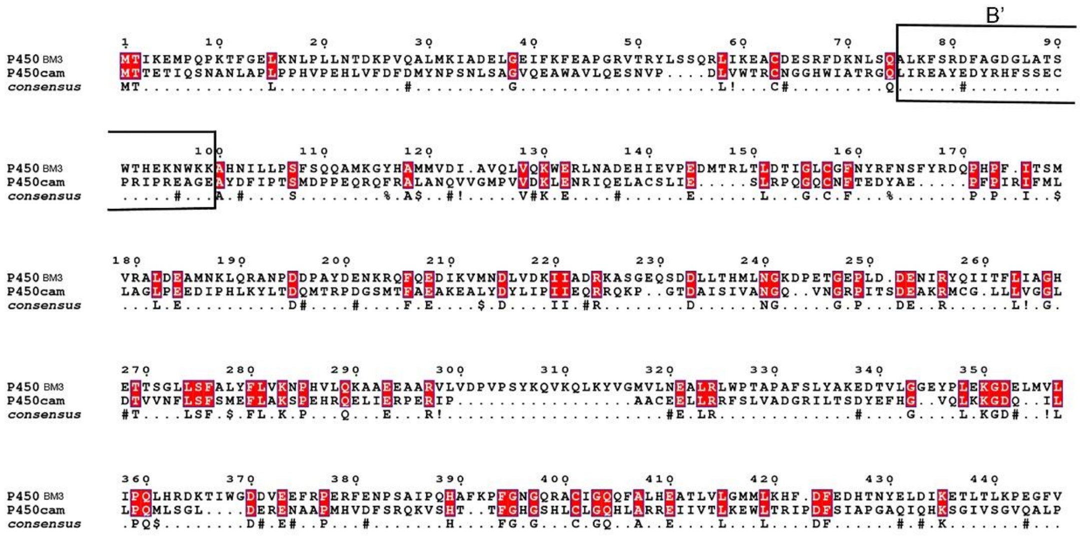

**Figure S3.** Protein sequence alignment of P450 BM3 and P450cam. Sequence alignment of P450 BM3 and P450cam based on their tertiary structures using T-COFFEE online service[16-17] and the figure was output by ESPript 3.0[18].

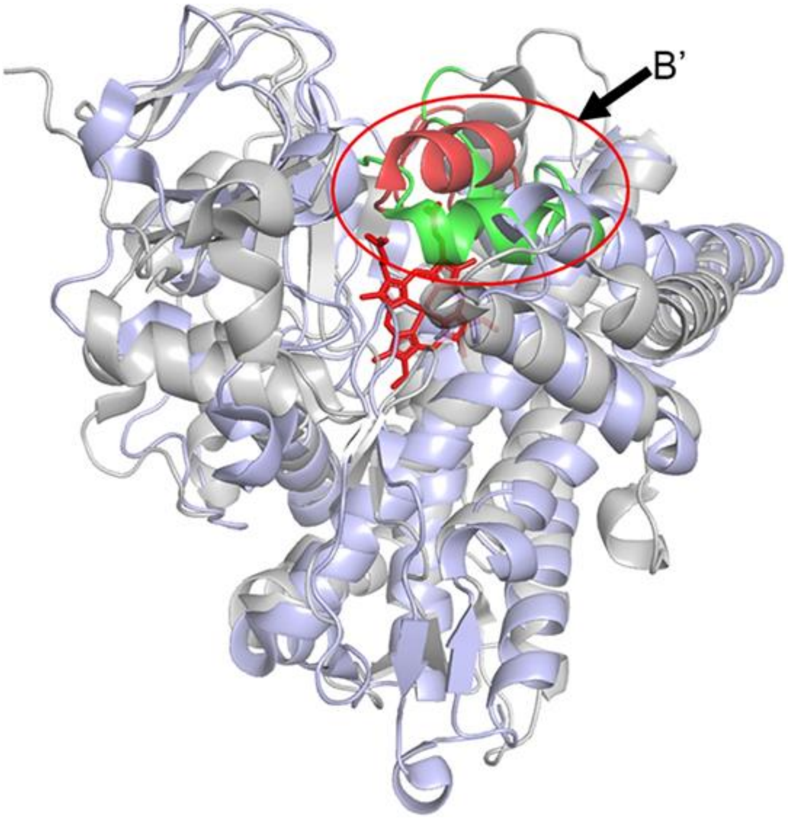

**Figure S4.** Structural comparison of P450 BM3 and P450cam. The superimposition of the three dimensional structures P450cam (PDB ID #: 2ZWU)[19] and P450 BM3 (PDB ID #: 2X80)[20]. The whole structures of P450cam and P450 BM3 are shown as cartoon in gray and light blue and their B’ helices colored in red and green respectively and marked by red ellipse.

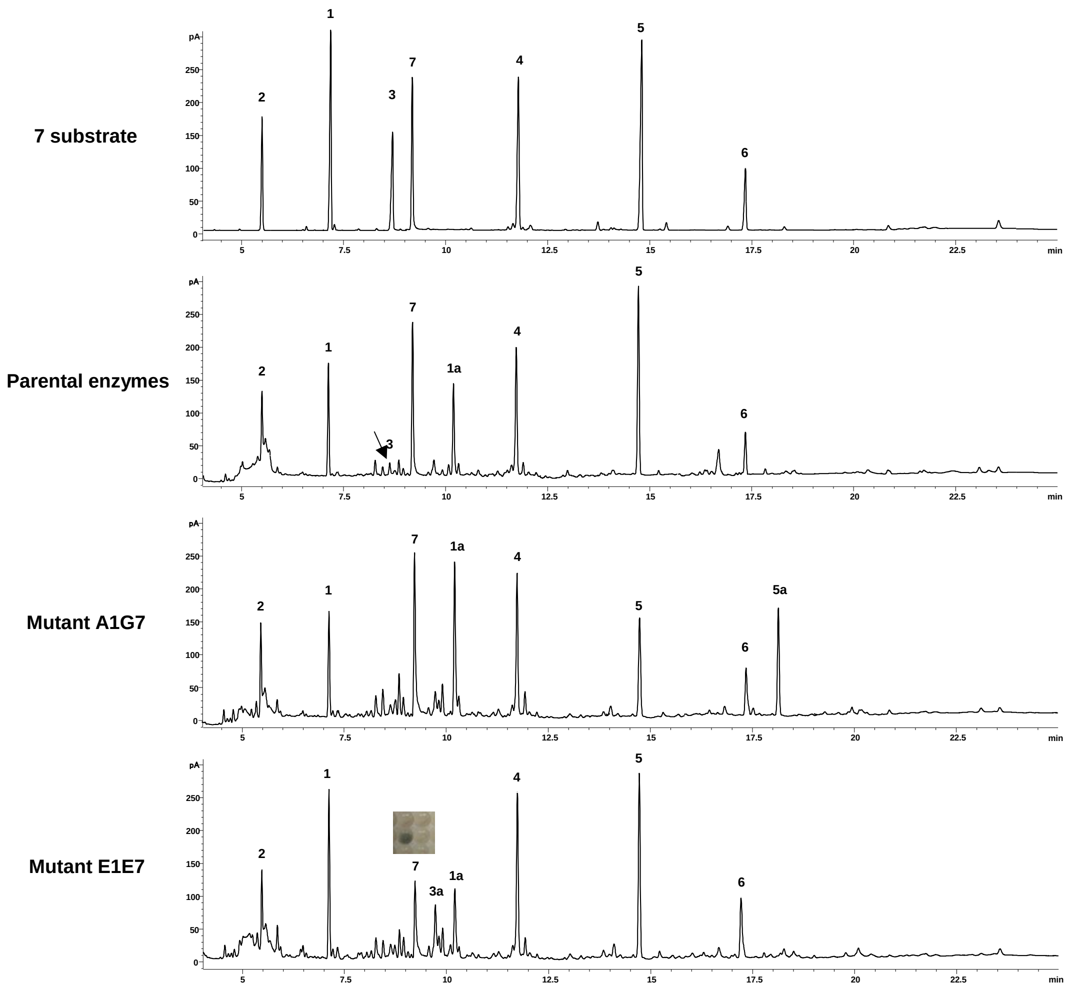

**Figure S5.** Representative GC chromatograms of the hit reactions. Compared with the reactions catalyzed by the parental P450 BM3 and P450cam, in the hit mixed reactions, the substrate peaks decreased and the product peaks were detected. The product of **7** was visible in blue color but not detectable by GC.

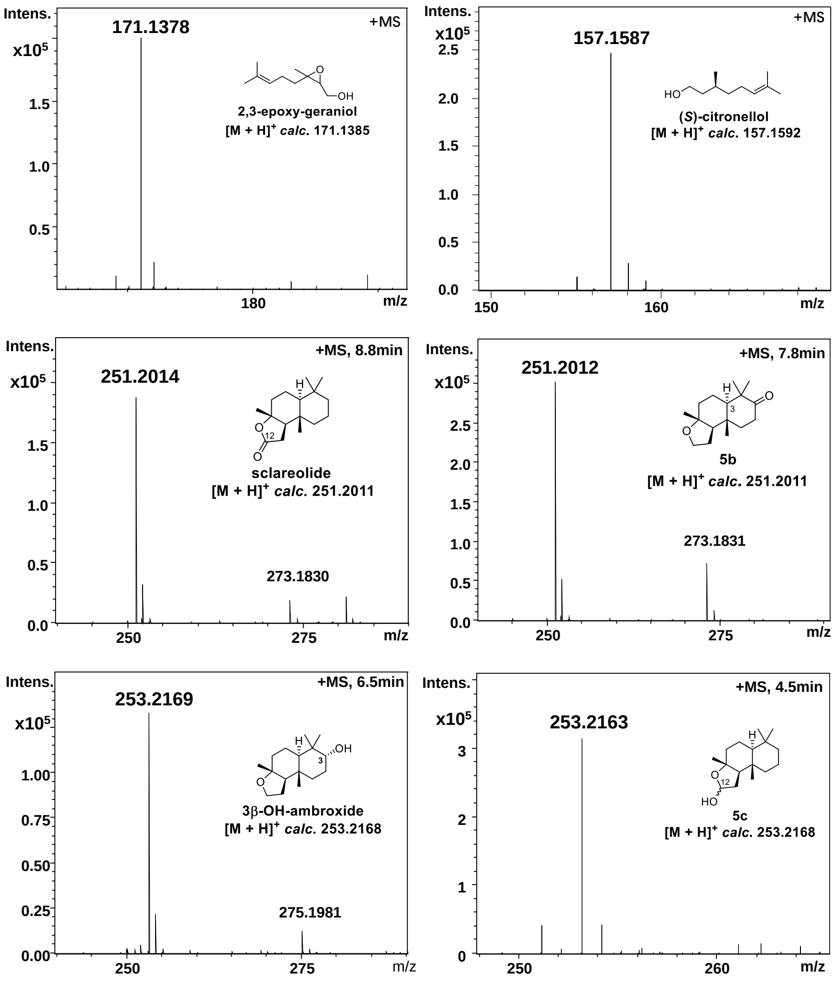

**Figure S6.** The high resolution mass spectra of products.

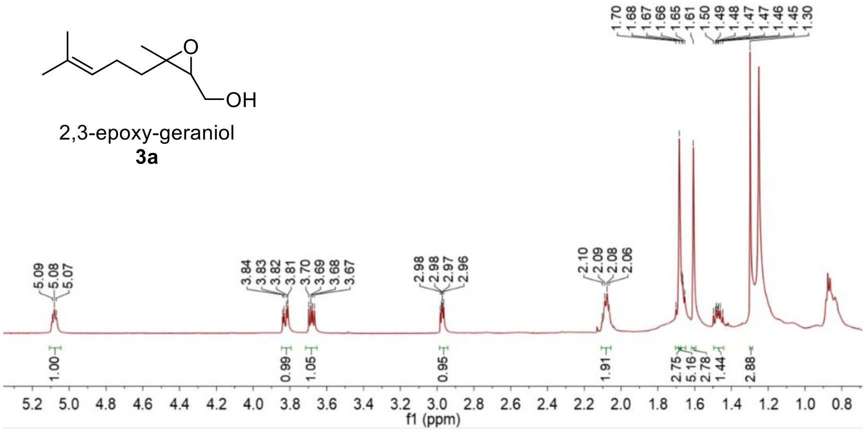

**Figure S7.** $^{1}$H NMR (500 MHz) spectrum of **3a** in $CDCl_3$.

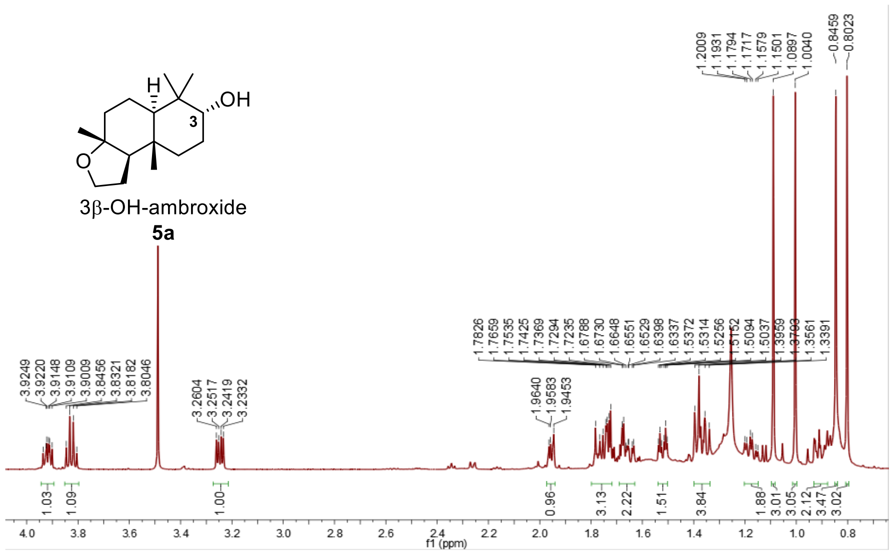

**Figure S8.** [1]H NMR (500 MHz) spectrum of **5a** in $CDCl_3$.

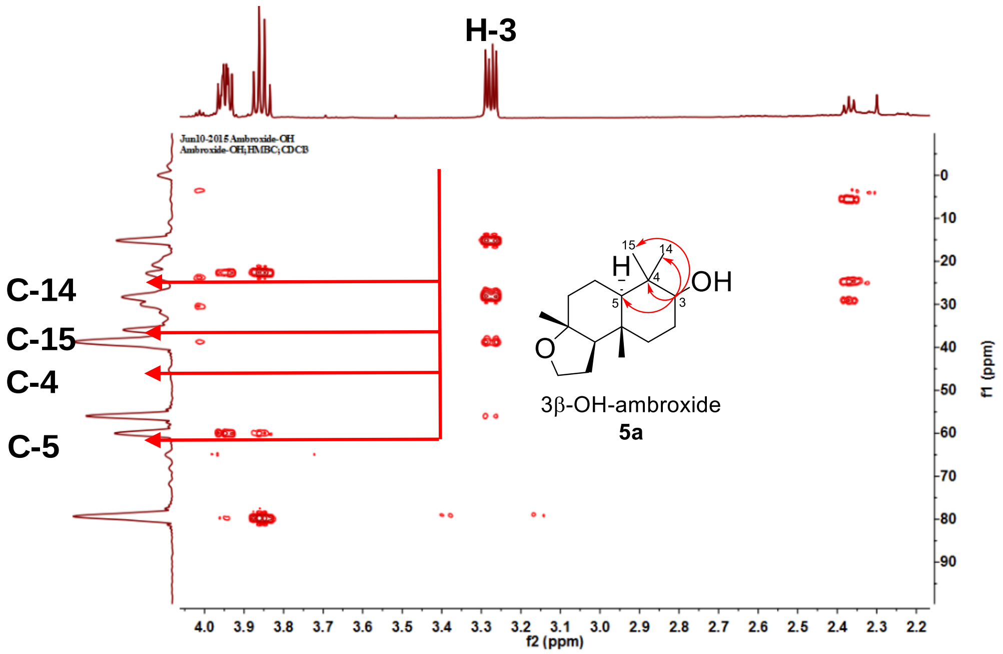

**Figure S9.** HMBC spectrum of **5a** in $CDCl_3$.

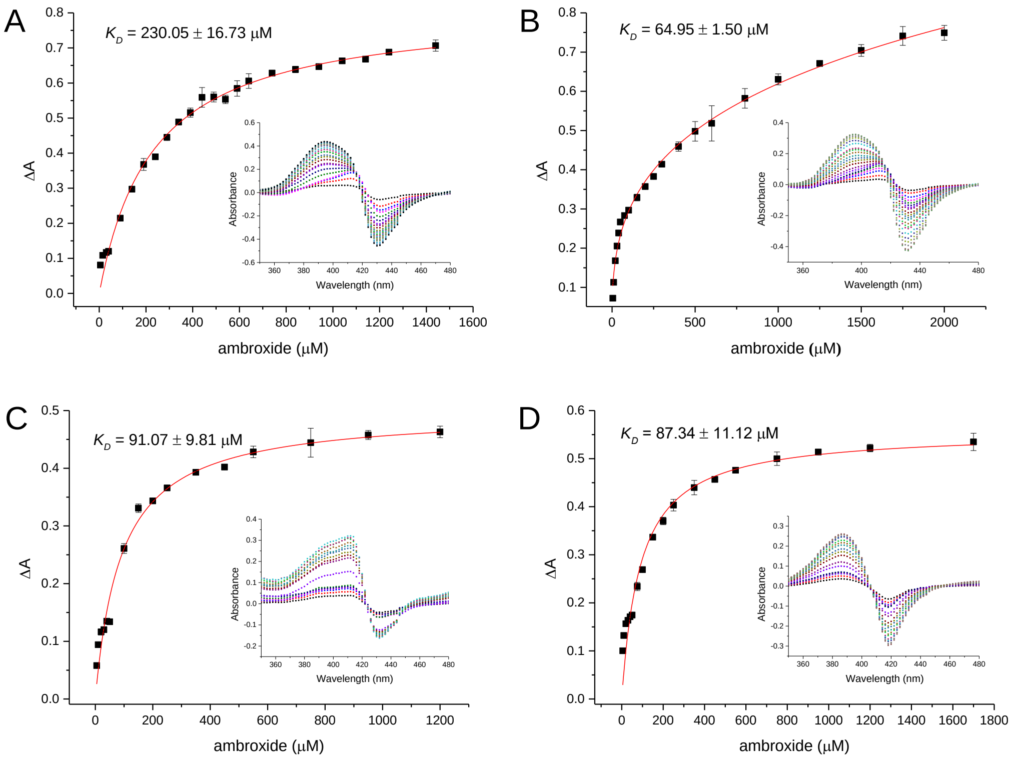

**Figure S10.** The substrate binding curves of **5** towards P450 BM3 (**A**), P450 BM3 V78S (**B**), P450cam (**C**), and P450cam F87R (**D**). The insets show the Type I binding spectra. All experiments were performed in duplicate.

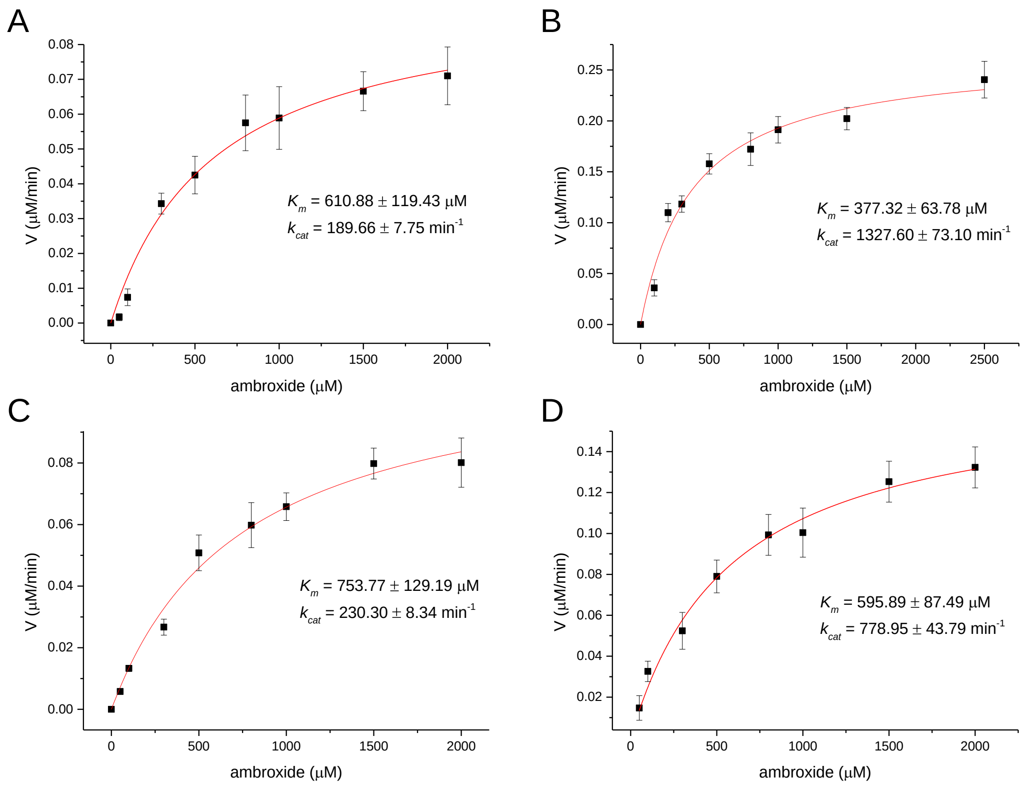

**Figure S11.** The kinetic curves of P450 BM3 (**A**), P450BM3 V78S (**B**), P450cam (**C**), and P450cam F87R (**D**) using **5** as substrate.

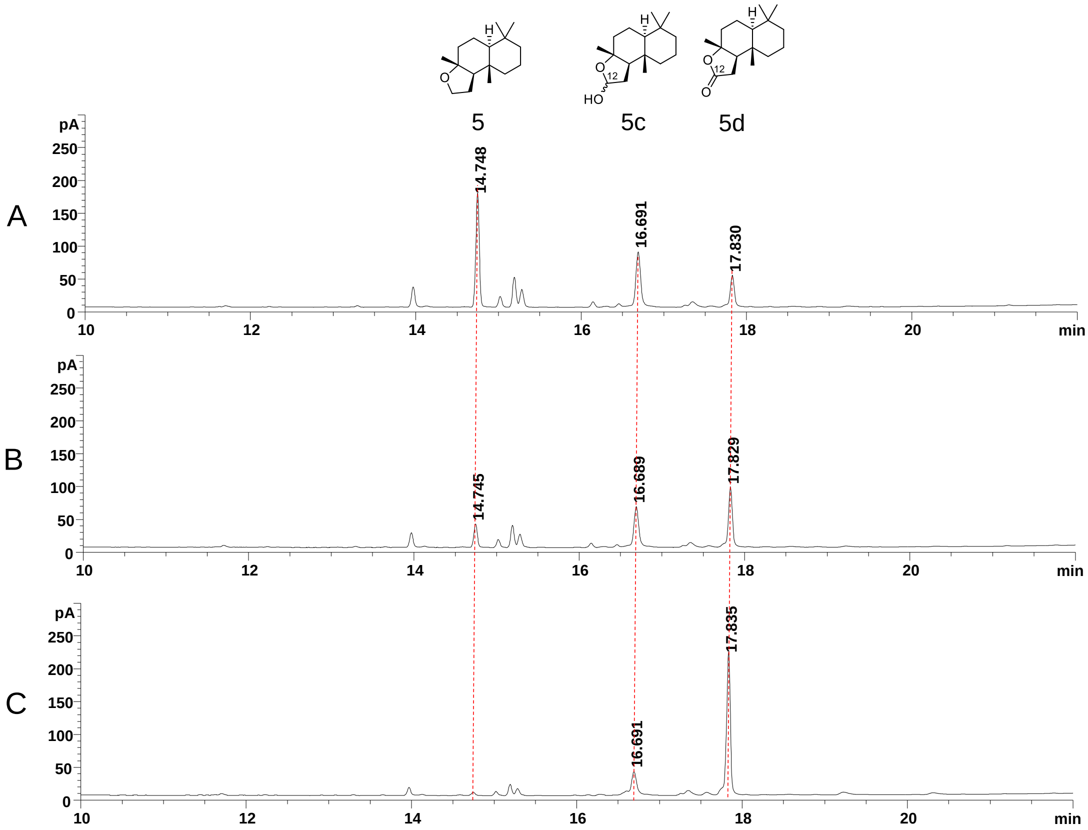

**Figure S12.** GC analysis of the product formation by P450 BM3 V78G. (A) 30 min; (B) 2 h; and (C) 3 h.

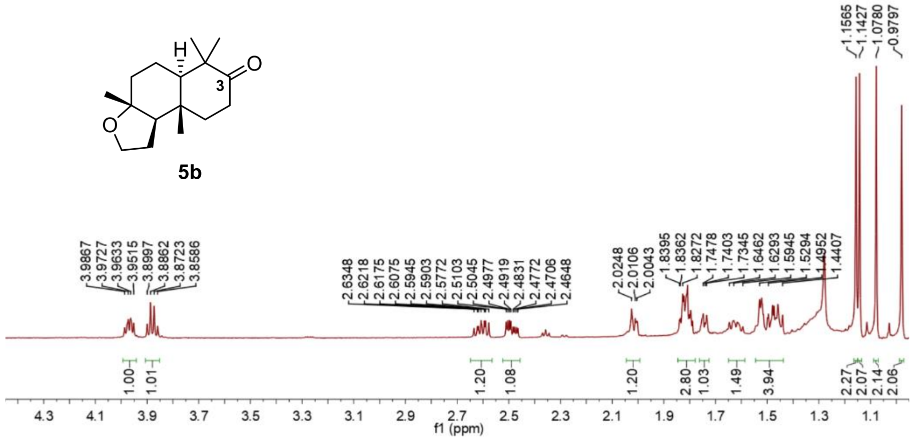

**Figure S13.** $^{1}H$ NMR (500 MHz) spectrum of **5b** in $CDCl_3$.

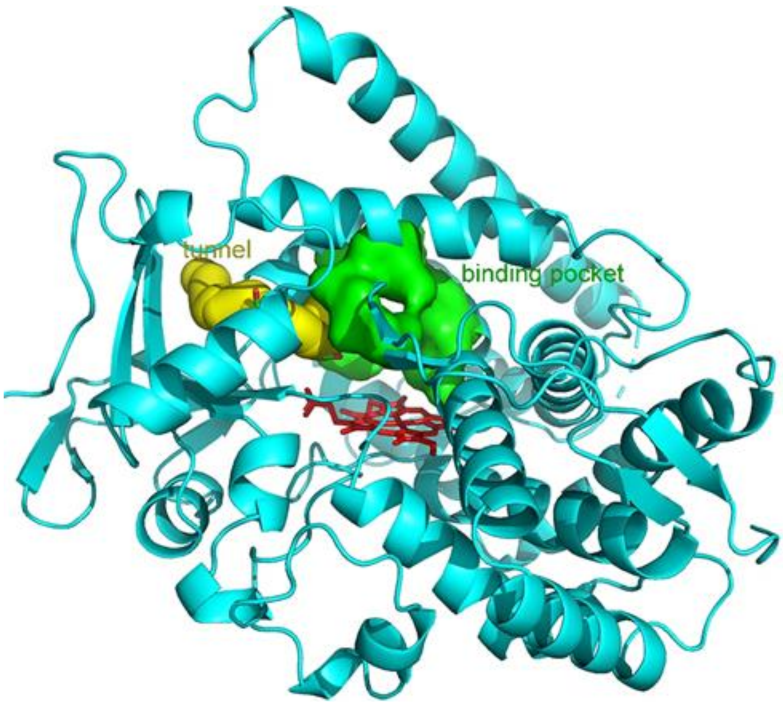

**Figure S14.** Structure of P450 BM3 V78S. The whole structure is shown as cartoon in cyan. The substrate binding pocket and the tunnel connecting the substrate binding pocket to outside solvent are shown as surface in green and yellow, respectively.

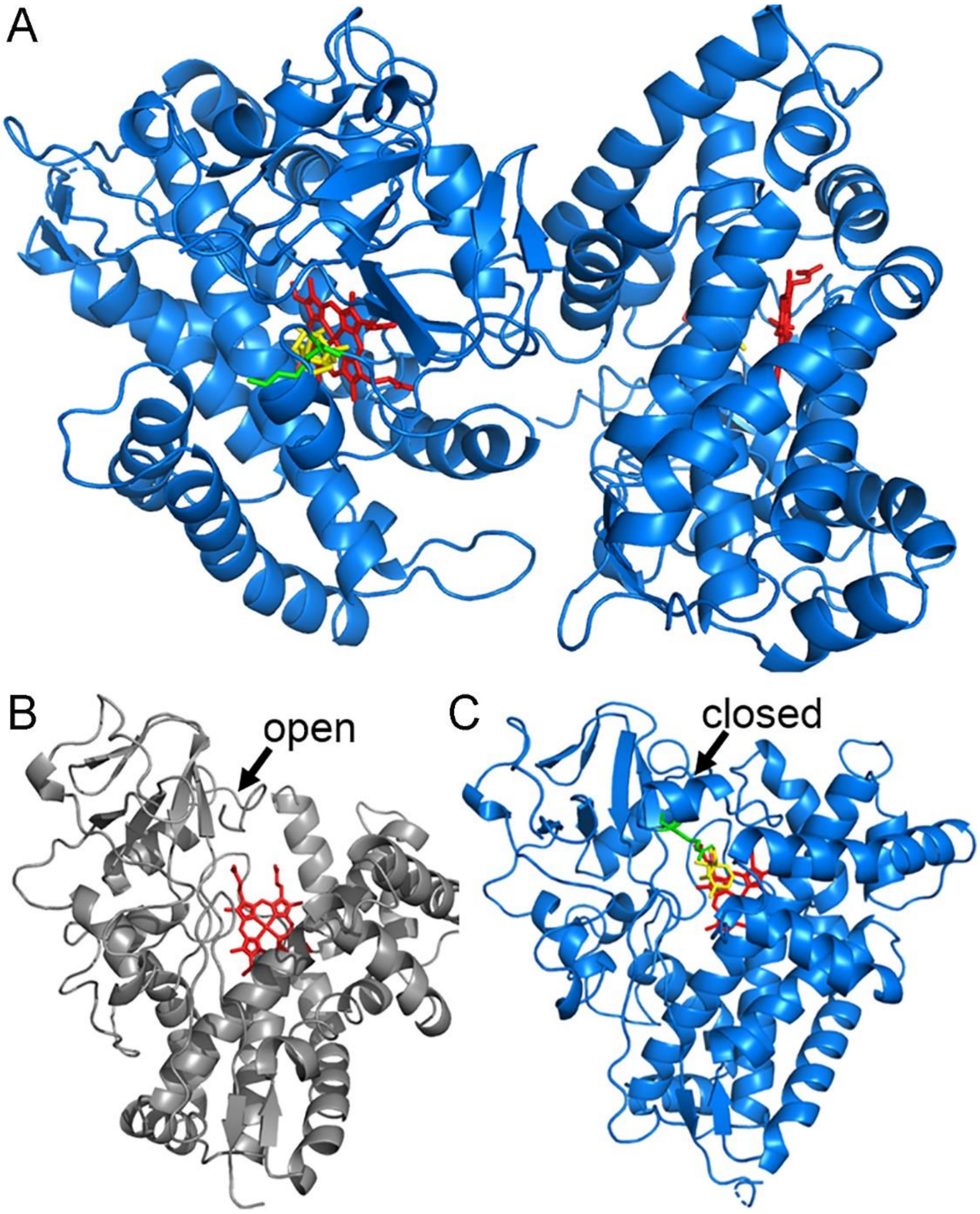

**Figure S15.** Structures of P450cam F87R. (**A**) Structure of **5**-bound P450cam F87R, which has two typical P450 folds existing in an asymmetric unit. Heme, R87 and substrate **5** shown as sticks in red, green and yellow respectively. (**B**) Strucure of substrate-free P450cam F87R in open state. (**C**) Structure of **5**-bound P450cam F87R in closed state. Heme, R87 and substrate **5** are shown as sticks in red, green and yellow, respectively.

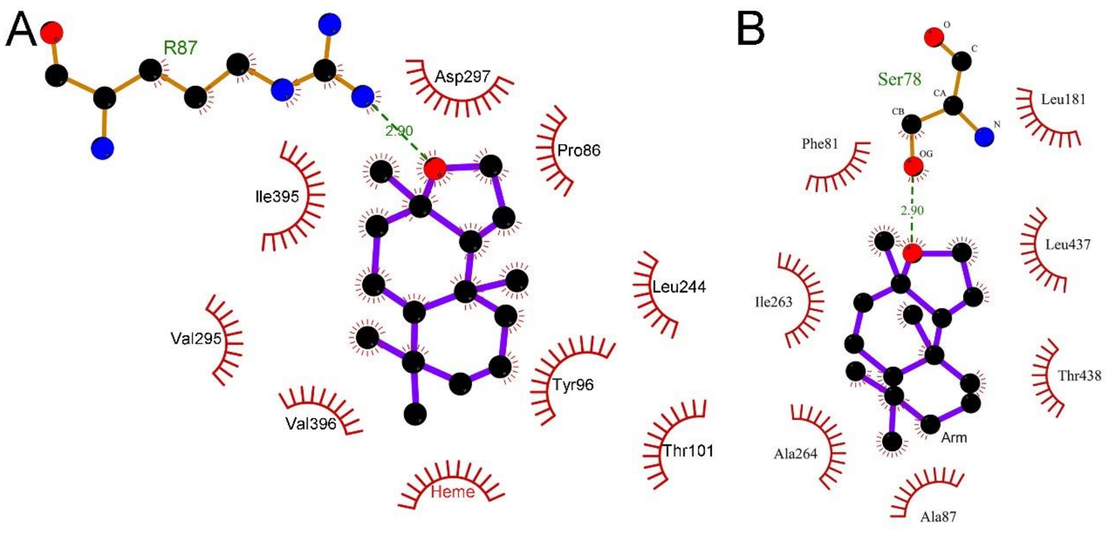

**Figure S16.** Key amino acids involved in substrate **5** binding. (**A**) Key amino acids around **5** in P450cam F87R complex structure; (**B**) Key amino acids around **5** in the ideal Autodock complex of P450 BM3 V78S. The figures are prepared using LigPlot+ (v.1.4.5).

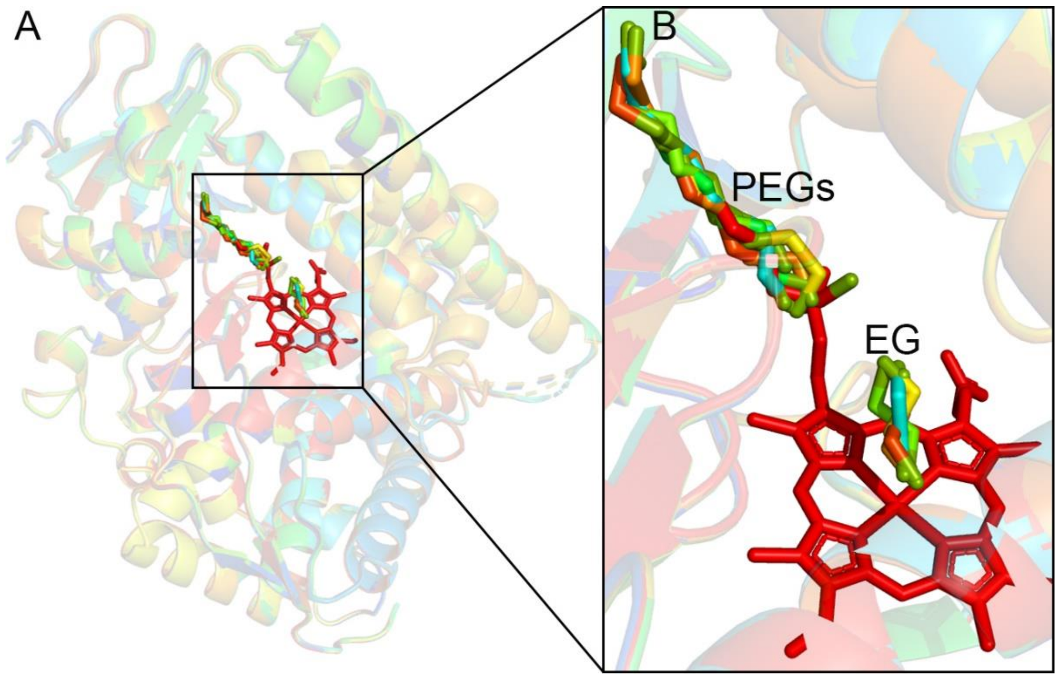

**Figure S17.** The overall structures of P450 BM3 V78S. (**A**) Superimposition of eight molecules of P450 BM3 V78S in an asymmetric unit. (**B**) PEGs with different carbon-chain length shown as sticks in different colors. EG (ethylene glycol) shown as sticks in different colors.

### References

(1) Zhang, W.; Du, L.; Li, F. W.; Zhang, X. W.; Qu, Z. P.; Hang, L.; Li, Z.; Sun, J. R.; Qi, F. X.; Yao, Q. P.; Sun, Y.; Geng, C.; Li, S. Y. Mechanistic Insights into Interactions between Bacterial Class I P450 Enzymes and Redox Partners. *ACS Catal.* **2018,** *8*, 9992-10003.
(2) Xue, Y. Q.; Wilson, D.; Zhao, L. S.; Liu, H. W.; Sherman, D. H. Hydroxylation of macrolactones YC-17 and narbomycin is mediated by the *pikC*-encoded cytochrome P450 in *Streptomyces venezuelae*. *Chem. Biol.* **1998,** *5*, 661-667.
(3) Du, L.; Dong, S.; Zhang, X.; Jiang, C.; Chen, J.; Yao, L.; Wang, X.; Wan, X.; Liu, X.; Wang, X.; Huang, S.; Cui, Q.; Feng, Y. Selective oxidation of aliphatic C-H bonds in alkylphenols by a chemomimetic biocatalytic system. *Proc. Natl. Acad. Sci. U. S. A.* **2017,** *114*, 5129-5137.
(4) Boddupalli, S. S.; Estabrook, R. W.; Peterson, J. A. Fatty acid monooxygenation by cytochrome P-450BM-3. *J. Biol. Chem.* **1990,** *265*, 4233-4239.
(5) Poulos, T. L.; Finzel, B. C.; Howard, A. J. High-resolution crystal structure of cytochrome P450cam. *J. Mol. Biol.* **1987,** *195*, 687-700.
(6) Guengerich, F. P.; Martin, M. V.; Sohl, C. D.; Cheng, Q. Measurement of cytochrome P450 and NADPH-cytochrome P450 reductase. *Nat. Protoc.* **2009,** *4*, 1245-1251.
(7) Gunsalus, I. C.; Wagner, G. C. Bacterial P-450cam methylene monooxygenase components: cytochrome m, putidaredoxin, and putidaredoxin reductase. *Methods Enzymol.* **1978,** *52*, 166-188.
(8) Li, S.; Podust, L. M.; Sherman, D. H. Engineering and analysis of a self-sufficient biosynthetic cytochrome P450 PikC fused to the RhFRED reductase domain. *J. Am. Chem. Soc.* **2007,** *129*, 12940-12941.
(9) Egami, H.; Oguma, T.; Katsuki, T. Oxidation Catalysis of Nb(salan) Complexes: Asymmetric Epoxidation of Allylic Alcohols Using Aqueous Hydrogen Peroxide as an Oxidant. *J. Am. Chem. Soc.* **2010,** *132*, 5886-5895.
(10) García-Granados, A.; Martínez, A.; Quirós, R.; Extremera, A. L. Chemical-microbiological semisynthesis of *enantio*-Ambrox® derivatives. *Tetrahedron* **1999,** *55*, 8567-8578.
(11) Otwinowski, Z.; Minor, W. Processing of X-ray diffraction data collected in oscillation mode. *Methods Enzymol.* **1997,** *276*, 307-326.
(12) McCoy, A. J.; Grosse-Kunstleve, R. W.; Adams, P. D.; Winn, M. D.; Storoni, L. C.; Read, R. J. Phaser crystallographic software. *J. Appl. Crystallogr.* **2007,** *40*, 658-674.
(13) Murshudov, G. N.; Vagin, A. A.; Dodson, E. J. Refinement of macromolecular structures by the maximum-likelihood method. *Acta Crystallogr., Sect. D: Biol. Crystallogr.* **1997,** *53*, 240-255.
(14) Emsley, P.; Lohkamp, B.; Scott, W. G.; Cowtan, K. Features and development of Coot. *Acta Crystallogr., Sect. D: Biol. Crystallogr.* **2010,** *66*, 486-501.
(15) Morris, G. M.; Huey, R.; Lindstrom, W.; Sanner, M. F.; Belew, R. K.; Goodsell, D. S.; Olson, A. J. AutoDock4 and AutoDockTools4: Automated docking with selective receptor flexibility. *J. Comput. Chem.* **2009,** *30*, 2785-2791.
(16) Notredame, C.; Higgins, D. G.; Heringa, J. T-Coffee: A novel method for fast and accurate multiple sequence alignment. *J. Mol. Biol.* **2000,** *302*, 205-217.
(17) Tommaso, P.; Moretti, S.; Xenarios, I.; Orobitg, M.; Montanyola, A.; Chang, J. M.; Taly, J. F.; Notredame, C. T-Coffee: a web server for the multiple sequence alignment of protein and RNA sequences using structural information and homology extension. *Nucleic Acids Res.* **2011,** *39*, 13-17.
(18) Robert, X.; Gouet, P. Deciphering key features in protein structures with the new ENDscript server. *Nucleic Acids Res.* **2014,** *42*, 320-324.
(19) Sakurai, K.; Shimada, H.; Hayashi, T.; Tsukihara, T. Substrate binding induces structural changes in cytochrome P450cam. *Acta Crystallogr. Sect. F: Struct. Biol. Cryst. Commun.* **2009,** *65*, 80-83.
(20) Kuper, J.; Tee, K. L.; Wilmanns, M.; Roccatano, D.; Schwaneberg, U.; Wong, T. S. The role of active-site Phe87 in modulating the organic co-solvent tolerance of cytochrome P450 BM3 monooxygenase. *Acta Crystallogr. Sect. F: Struct. Biol. Cryst. Commun.* **2012,** *68*, 1013-1017.
